# Cells use molecular working memory to navigate in changing chemoattractant fields

**DOI:** 10.1101/2021.11.11.468222

**Authors:** Akhilesh Nandan, Abhishek Das, Robert Lott, Aneta Koseska

## Abstract

In order to migrate over large distances, cells within tissues and organisms rely on sensing local gradient cues. These cues however are multifarious, irregular or conflicting, changing both in time and space. Here we find that single cells utilize a molecular mechanism akin to a working memory, to generate persistent directional migration when signals are disrupted by temporally memorizing their position, while still remaining adaptive to spatial and temporal changes of the signal source. Using dynamical systems theory, we derive that these information processing capabilities are inherent for protein networks whose dynamics is maintained away from steady state through organization at criticality. We demonstrate experimentally using the Epidermal growth factor receptor (EGFR) signaling network, that the memory is maintained in the prolonged receptor’s activity via a slow-escaping remnant, a dynamical ”ghost” of the attractor of the polarized signaling state, that further results in memory in migration. As this state is metastable, it also enables continuous adaptation of the migration direction when the signals vary in space and time. We therefore show that cells implement real-time computations without stable-states to navigate in changing chemoattractant fields by memorizing position of disrupted signals while maintaining sensitivity to novel chemical cues.

## Introduction

Directed chemotactic behavior relies on generating polarized signaling activity at the plasma membrane of the cell that is translated to an elongated cell shape in the direction of the signal. Cells maintain the acquired orientation longer than the duration of the recently encountered signal in order to avoid immediate switching to random migration when signals are temporarily disrupted or noisy, while simultaneously remaining sensitive and are able to adapt the migration direction based on the changes in the environment (Parent and Devreotes, 1999; Foxman et al., 1999; Ridley et al., 2003). Thus, cells as diverse as social amoeba, neutrophils, leukocytes, fibroblasts and nerve cells, not only respond to dynamic gradients, but also integrate and resolve competing spatial signals or prioritize newly encountering attractants, likely by memorizing their recent environment (Jilkine and Edelstein-Keshet, 2011; Skoge et al., 2014; Albrecht and Petty, 1998). Numerous models of chemotactic responses based on positive feedbacks, incoherent feed-forward, excitable or Turing-like networks have been proposed, accounting either for sensing non-stationary stimuli or for long-term maintenance of polarized signaling activity, but not both (Levchenko and Iglesias, 2002; Levine et al., 2002; Mori et al., 2008; Goryachev and Pokhilko, 2008; Beta et al., 2008; Xiong et al., 2010; Trong et al., 2014; Halatek and Frey, 2018). These models rely on computations with stable states, where switching from the attractor of basal- to the attractor of polarized-signaling activity enables noise-robust sensing, or establishing a long-term memory of previous signal localization. However, they are less suited for real-time computation of signals that vary in time and space, since the stable attractors completely hinder or at least significantly delay the responsiveness to newly encountered signals (Stanoev et al., 2020). Thus, the mechanism that underlies robust cellular navigation in changing chemical fields has remained unknown.

Here we set out to identify how cells satisfy these two general, but seemingly opposed demands: maintaining temporal memory in directional migration through a prolonged polarized state beyond the chemotactic signal duration, while still being able to quickly reset and re-adapt upon novel sensory cues. Using a mathematical model of EGFR network signaling dynamics, we predict and demonstrate experimentally in epithelial cells that these competing demands are uniquely fulfilled for network’s organization at criticality. Beyond this specific biological implementation, we present a generic dynamical mechanism that addresses how cells compare and integrate chemical cues over time and space in order to generate robust responses in a history-dependent manner.

## Results

### 1 Dynamical basis of navigation in non-stationary environments

We conjectured that operating in changing environments likely relies on computations with metastable states rather than stable attractors, to allow both for transient stability of the polarized signaling state when signals are disrupted or noisy, as well as its rapid adaptation when the signals vary in space and time. Our hypothesis is that this can be achieved if biochemical systems are maintained away from steady state. We therefore approached the problem using the abstract language of dynamical systems theory, where the characteristics of any process directly follow from the type of dynamical transitions, called bifurcations, through which they emerge (Strogatz, 2018). In our previous work we identified that when a saddle-node bifurcation (*SN*) and thereby a steady-state is lost in a dynamical transition, i.e. upon signal removal, a remnant or a dynamical ”ghost” of the stable attractor serves as a mechanism for sensing timevarying growth factors in biochemical receptor networks (Stanoev et al., 2018; Stanoev et al., 2020). Necessary for manifestation of the ”ghost” state is organization at criticality, which in the networks we previously examined was determined by the concentration of receptors on the cell membrane. Moreover, the ”ghost” state is dynamically metastable and transiently maintains the system in the vicinity of the steady state. Thus, a transient memory of previously present stimuli is generated, which enables integration of information contained in the temporal signals. Navigation in changing environments through directed migration however, must additionally rely on a polarized representation of the directional signal, requiring a reliable mechanism for signal-induced transition from a non-polarized symmetric to an asymmetric polarized receptor signaling state and subsequently polarized cell shape. From a dynamical systems point of view, a pitchfork bifurcation (*PB*) can satisfy the condition for robust cell polarization, since by definition a *PB* characterizes a transition from a homogeneous to an inhomogeneous steady state (Koseska et al., 2013; Strogatz, 2018). We thus hypothesized that organization at criticality - in the vicinity of a *SN*_*PB*_ through which a sub-critical *PB* is stabilized (grey shaded area in Figure 1A), the dynamical characteristics of both bifurcations, signal integration through dynamic memory and cell polarization, will be uniquely manifested to render a minimal mechanism for responsiveness in changing environments.

**Figure 1.**
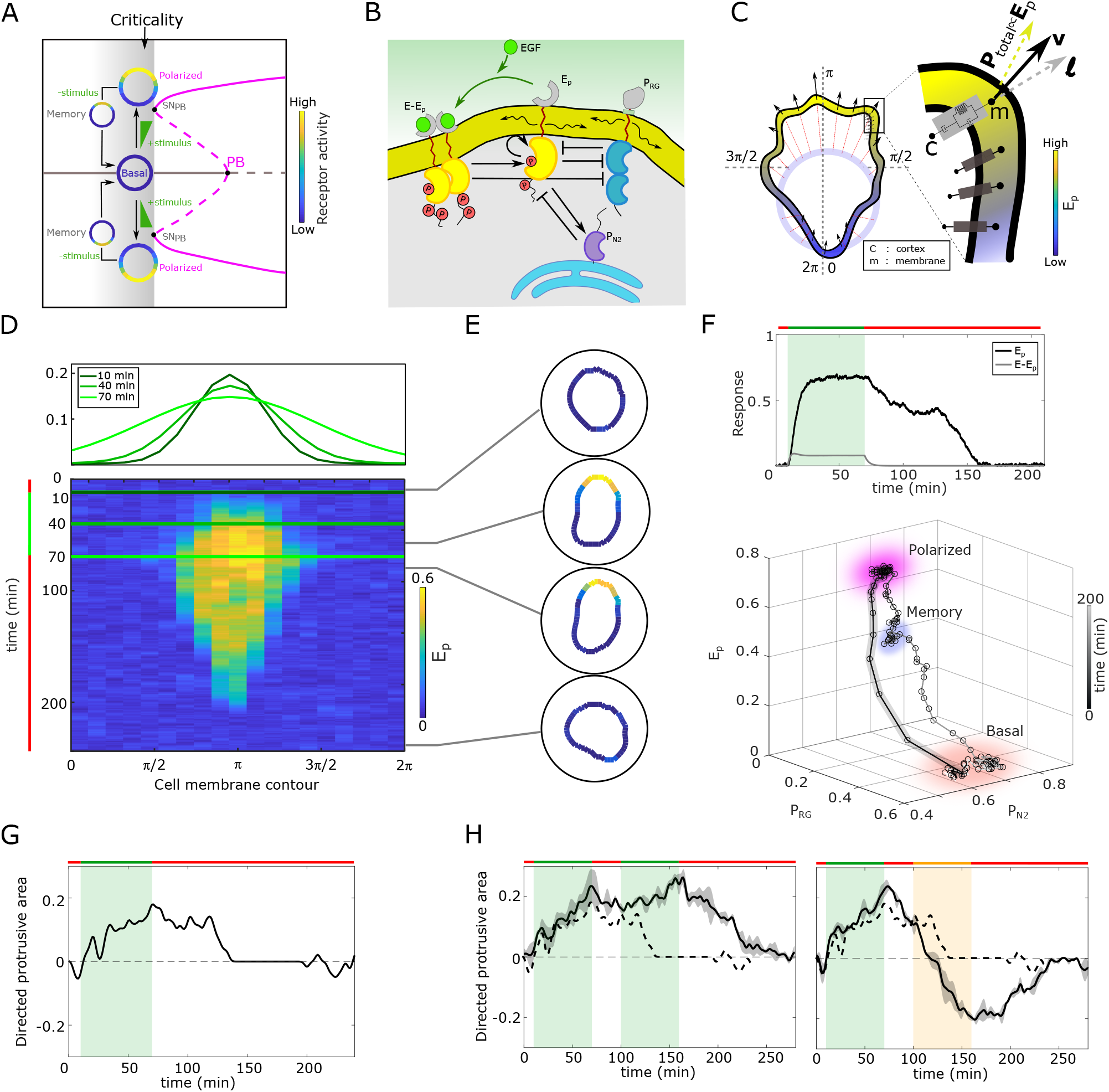
Organization at criticality enables sensing changing spatial-temporal signals. **A**, Dynamical mechanism: critical organization before sub-critical pitchfork bifurcation (*PB*, grey shaded area). Stable/unstable steady states (solid/dashed lines): basal (homogeneous, grey) and polarized (inhomogeneous, magenta) receptor activity; stimulus induced transitions between states: arrow lines. *SN*_*PB*_: saddle-node bifurcation through which *PB* is stabilized. **B**, Scheme of the EGFR-PTP interaction network. Ligandless EGFR (*E*_*p*_) interacts with PT-PRG (*P*_*RG*_) and PTPN2 (*P*_*N*2_). Liganded EGFR (*E* − *E*_*p*_) promotes autocatalysis of *E*_*p*_. Causal links: solid black lines; curved arrow lines: diffusion. See also Figure S1A. **C**, Signal-induced shape-changes during cell polarization. Arrows: local edge velocity direction. Zoom: Viscoelastic model of the cell - parallel connection of an elastic and a viscous element. **P**_**total**_: total pressure; **v**: local membrane velocity; **l**: viscoelastic state. Bold letters: vectors. Cell membrane contour: [0, 2*π*]. **D**, Top: *In silico* evolution of spatial EGF distribution. Bottom: Kymograph of *E*_*p*_ for organization at criticality from reaction-diffusion simulations of the network in (**B**). **E**, Corresponding exemplary cell shapes with color coded *E*_*p*_, obtained with the model in (**C**). **F**, Top: Temporal profiles *E*_*p*_ (black) and *E* − *E*_*p*_ (grey). Green shaded area: EGF gradient presence. Bottom: State-space trajectory of the system with denoted trapping state-space areas (colored). See also movie S1. Thick/thin line: signal presence/absence. **G**, Quantification of *in silico* cell morphological changes from the example in **E. H**, Left: same as in **G**, only when stimulated with two consecutive dynamic gradients from same direction. Second gradient within the memory phase of the first. Right: the second gradient (orange) has opposite localization. Mean±s.d. from n=3. See also Figure S1C,D. Dashed line: curve from **G**. Parameters: Supplementary information. In **(D-H)**, green(orange)/red lines: stimulus presence/absence. See also Figure S1.

We described this conjecture mathematically for a general reaction-diffusion model representing the signaling activity on the plasma membrane of a cell, 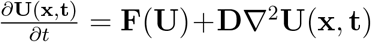, with **U** being the vector of local densities of active signaling components, **D** - diffusion constants and **F** accounting for all chemical reactions. Our analysis shows that a *PB* exists if, for a spatial perturbation of the symmetric steady state (**U**_**s**_) of the form **U**(**x**, *t*) = **U**_**s**_ + *δ***U**(**x**)*e*^*λt*^, the conditions *δ***U**(−**x**) = −*δ***U**(**x**) and the limit lim_*λ*→0_ *F*_*λ*_ = *det*(*J*) = 0 are simultaneously fulfilled (Supplementary information). This implies that the linearized system has zero-crossing eigenvalues (*λ*) associated with the odd mode of the perturbation (Paquin-Lefebvre et al., 2020). To probe the sub-critical transition and therefore the necessary organization at criticality, a reduced description in terms of an asymptotic expansion of the amplitude of the polarized state (*φ*) must yield the Landau equation 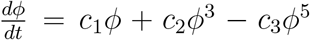, guaranteeing the existence of *SN*_*PB*_ (see Supplementary information for derivation).

These abstract dynamical transitions can be realized in receptor tyrosine kinase signaling networks with different topologies and are best analyzed using computational models, whose predictions are then tested in quantitative experiments on living cells. To exemplify the above mentioned principle, we use the well-characterized Epidermal growth factor receptor (EGFR) sensing network (Reynolds et al., 2003; Baumdick et al., 2015; Stanoev et al., 2018). It constitutes of double negative and negative feedback interactions of the receptor, EGFR (*E*_*p*_) with two enzymes, the phosphatases PTPRG (*P*_*RG*_) and PTPN2 (*P*_*N*2_; Figure 1B, Figure S1A), respectively. *E*_*p*_ and *P*_*RG*_ laterally diffuse on the membrane and inhibit each-other’s activities (see Supplementary information for the molecular details of the network). These molecular interactions can be mathematically described using mass action kinetics (Eqs.(6) in Supplementary information), and a weakly nonlinear analysis (Becherer et al., 2009) shows that the EGFR signaling dynamics undergoes a symmetry-breaking transition as outlined above (proof in Supplementary information, Figure S1B). Contrary to a bistable system, where the polarized signaling state would be manifested by two steady states, for e.g. high and low protein phosphorylation in the front and back of the cell respectively (Beta et al., 2008), the inhomogeneous steady state generated via a *PB* is a single attractor defined as a combination of the front and back activity states. This profiles *PB* as a robust mechanism of cell polarization.

Polarized EGFR signaling on the other hand, will lead to reorganization of the cortical actomyosin cytoskeleton by regulating members of the Rho GTPase family, thereby inducing signal-dependent cell shape changes and subsequent migration (Chiasson-MacKenize and McClatchey, 2018; Ridley and Hall, 1992). In order to link signaling activity with morphodynamics, we modeled the cell as a viscoelastic cortex surrounding a viscous core (Yang et al., 2008) (Supplementary information), where EGFR signaling dynamics affects cell shape changes through the protrusion/retraction stress and the viscoelastic nature of the cell membrane (Figure 1C).

We first fixed the total EGFR concentration on the cell membrane to a value that corresponds to organization at criticality, and investigated the response of the *in silico* cell to gradient stimulus. In the absence of stimulus, EGFR phosphorylation is uniformly distributed along the cell membrane rendering a symmetrical cell shape (Figure 1D, E). Introducing dynamic gradient stimulus in the simulation (slope changes from steep to shallow over time, Figure 1D, top) led to rapid polarization of EGFR phosphorylation in the direction of the maximal chemoattractant concentration, generating a cell shape with a clear front and back. The polarized signaling state was maintained for a transient period of time after removal of the gradient, corresponding to manifestation of memory of the localization of the previously encountered signal (Figures 1D,E; temporal profile Figure 1F, top). The prolonged polarized state does not result from remnant ligand-bound receptors (*E* − *E*_*p*_) on the plasma membrane, as they exponentially decline after signal removal (Figure 1F, top). The memory in polarized signaling was also reflected on the level of the cell morphology, as shown by the difference of normalized cell protrusion area in the front and the back of the cell over time (Figure 1G). Plotting the trajectory that describes the change of the state of the system over time (state-space trajectory, Figure 1F bottom, movie S1) shows that the temporal memory in EGFR phosphorylation polarization is established due to transient trapping of the signaling state trajectory in state-space. This is typical for the emergence of metastable ”ghost” states (Stanoev et al., 2020; Strogatz, 2018), indicating that the system is maintained away from steady-states. The trapping in the dynamically-metastable memory state does not hinder sensing of and adapting to subsequent signals. The cell polarity is sustained even when the EGF signal is briefly disrupted, and the cell is able to reverse direction of polarization when the signal direction is inverted (Figure 1H, Figures S1C,D). We next chose in the simulations a higher EGFR concentration on the membrane, such that the system moves from criticality to organization in the stable inhomogeneous state regime. In this scenario, even a transient signal induces switching to the polarized state that is permanently maintained, generating a long-term memory of the direction on the initial signal. Thus, the cell is insensitive to subsequent stimuli from the same direction, whereas consecutive gradients from opposite directions generate conflicting information that cannot be resolved (Figure S1E). Organization in the homogeneous, symmetric steady states on the other hand renders cells insensitive to the extracellular signals (Figure S1F,G). These response features for organization in the stable steady state regimes resemble the finding of previously published models: such models cannot simultaneously capture memory in polarization along with continuous adaptation to novel signals, or require fine-tuning of kinetic parameters to explain the experimentally observed cell behavior (Levchenko and Iglesias, 2002; Levine et al., 2002; Mori et al., 2008; Goryachev and Pokhilko, 2008; Beta et al., 2008; Xiong et al., 2010; Trong et al., 2014). This demonstrates that organization at criticality, in a vicinity of a *SN*_*PB*_, is a unique mechanism for processing changing signals.

### 2 Cells display temporal memory in polarized receptor phosphorylation resulting from a dynamical ”ghost”

To test experimentally whether cells maintain memory of the direction of previously encountered signals and what is the duration of this effect, epithelial breast cancer-derived MCF7 cells were subjected for 1h to a stable gradient of fluorescently tagged EGF-Alexa647 (EGF^647^) with a maximal amplitude of 10ng/ml applied from the top of the chamber in a computer-programmable microfluidic device (Figures 2A,B). EGFR phosphorylation at the plasma membrane was quantified during and for 3h after gradient wash-out by determining the rapid translocation of mCherry-tagged phosphotyrosine-binding domain (PTB^*mCherry*^) to phosphorylated tyrosines 1086/1148 of ectopically expressed EGFR-mCitrine (EGFR^*mCitrine*^) using ratiometric imaging (Offterdinger et al., 2004)(Methods). Due to the low endogenous EGFR levels in MCF7 cells, the expression range of EGFR^*mCitrine*^ was set to mimic the endogenous receptor range in the related MCF10A cell line, such that both cell lines have equivalent signaling properties of downstream effector molecules (Stanoev et al., 2018), and were therefore used in a complementary way in this study.

**Figure 2.**
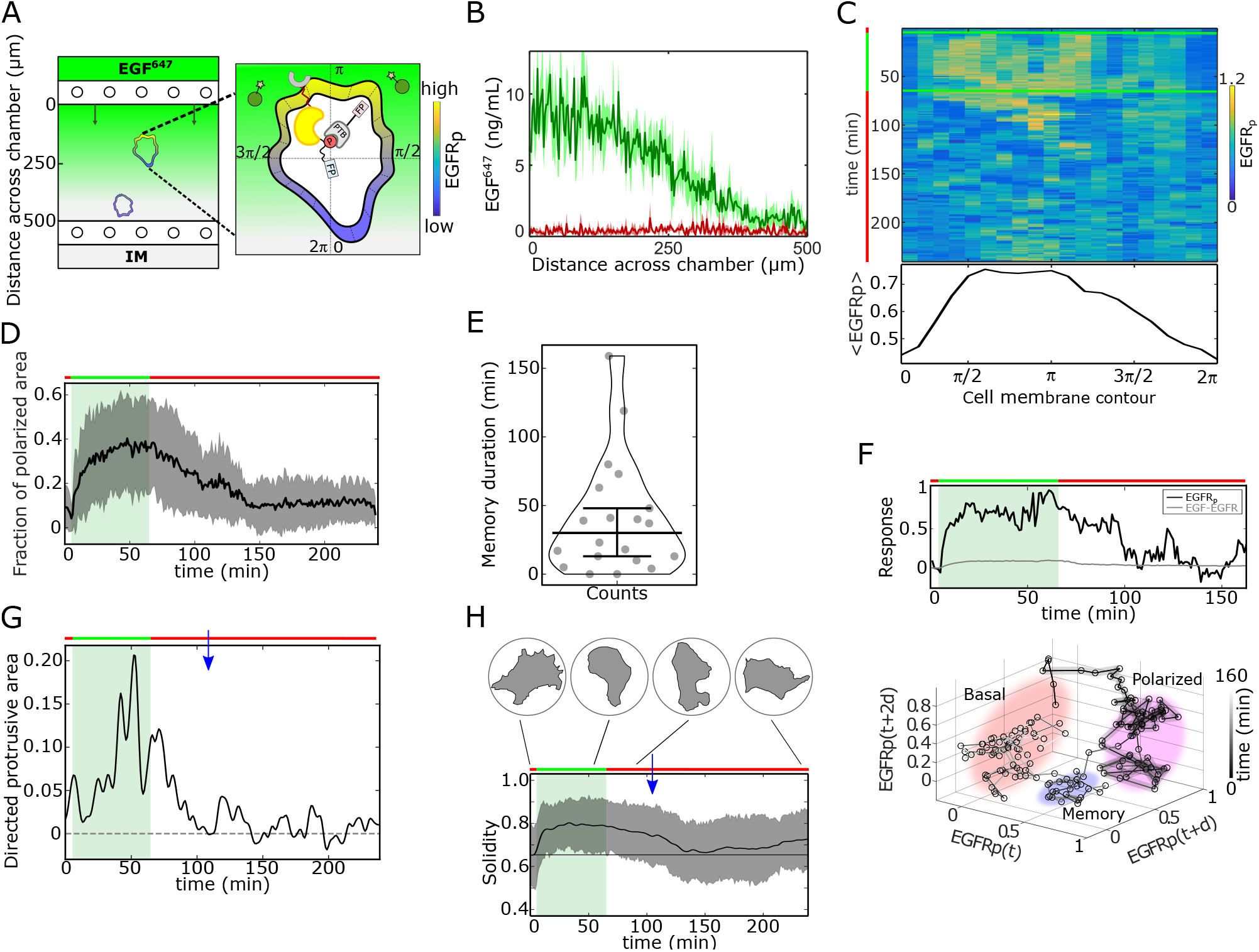
Single-cell molecular memory in polarized EGFR^*mCitrine*^ phosphorylation from dynamical state-space trapping. **A**, Scheme of microfluidic EGF^647^-gradient experiment; Zoom: single-cell measurables. Cell membrane contour [0, 2*π*] (20 segments). *PTB* - phosphotyrosine binding domain, *FP* /star symbol - fluorescent protein, *EGFR*_*p*_-phosphorylated EGFR^*mCitrine*^. Remaining symbols as in Figure 1B. **B**, Quantification of EGF^647^ gradient profile (at 60*min*, green) and after gradient wash-out (at 65*min*, red). Mean±s.d., N=4. **C**, Exemplary quantification of, Top: single-cell *EGFR*_*p*_ kymograph. Data was acquired at 1*min* intervals in live MCF7-EGFR^*mCitrine*^ cells subjected for 60*min* to an EGF^647^ gradient. Other examples in Figure S2D. Bottom: respective spatial projection of *EGFR*_*p*_. Average using a moving window of 7 bins is shown. Mean±s.d. from n=20, N=7 in Figure S2C. **D**, Average fraction of polarized plasma membrane area (mean±s.d.). In **D, E** and **H**, n=20, N=7. **E**, Quantification of memory duration in single cells (median±C.I.). **F**, Top: Temporal profiles of phosphorylated *EGFR*^*mCitrine*^ (black) and *EGF* ^647^ − *EGFR*^*mCitrine*^ (grey) corresponding to **C**. Bottom: Corresponding reconstructed state-space trajectory (movie S2) with denoted trapping state-space areas (colored). Thick/thin line: signal presence/absence. d - embedding time delay. **G**, Exemplary quantification of morphological changes, directed cell protrusion area, for the cell in **C**. Memory duration: 43*min*. **H**, Averaged single-cell morphological changes (Solidity, mean±s.d.). Average memory duration: 40*min*. Top insets: representative cell masks at distinct time points. In **D, F-H**, green shaded area: EGF^647^ gradient duration; green/red lines: stimulus presence/absence. Blue arrow: end of memory. See also Figure S2.

Kymograph analysis of EGFR^*mCitrine*^ phosphorylation at the plasma membrane of single cells showed polarization in gradient of EGF^647^ (Figure 2C, Figures. S2A-D), as shallow as 10% between front and back of the cell. Only few cells manifested basal or symmetric EGFR^*mCitrine*^ phosphorylation distribution upon gradient stimulation (Figures S2A-B,E). Quantifying the fraction of plasma membrane area with polarized EGFR^*mCitrine*^ phosphorylation revealed that the polarization persisted ∼ 40*min* on average after gradient removal ([4 − 159*min*] Figures 2D,E; Figure S2F). In order to identify whether the experimentally observed memory results from a dynamically metastable (transiently stable) signaling state, we next reconstructed the state-space trajectory from the measured single-cell temporal EGFR^*mCitrine*^ posphorylation profile using Takens’s delay embedding theorem (Takens, 1980)(Methods). Trajectory trapping in a state-space area different than that of the polarized and basal steady states characterized the memory phase, corroborating that the memory in EGFR^*mCitrine*^ phosphorylation polarization emerges from a *SN*_*PB*_ ”ghost” that maintains the system away from the steady-states (compare Figure 2F to 1F, movie S2). Although the memory doesn’t result from a stable state, it enables to maintain memory of the polarized cell morphology even after gradient removal. This is reflected through the exemplary temporal evolution of the cell protrusion area in direction of the gradient (Figure 2G, memory duration ∼ 43*min*). On average, single epithelial cells maintained the polarized cell shape ∼ 40*min* after signal removal (Figure 2H, Methods). The average duration of memory in the polarized cell morphology therefore directly corresponds to the average memory duration in signaling, suggesting that it will be also reflected as memory in directed cell migration.

### 3 Transient memory in cell polarization is translated to transient memory in directional migration

To test the phenotypic implications of the transient memory in cell polarization, we analyzed the motility features of the engineered MCF7-EGFR^*mCitrine*^, as well as of MCF10A cells at physiological EGF concentrations. Cells were subjected to a 5h dynamic EGF^647^ gradient that was linearly distributed within the chamber, with EGF^647^ ranging between 25−0ng/ml, allowing for optimal cell migration (Figure S3A). The gradient steepness was progressively decreased in a controlled manner, rendering an evolution towards a ∼ 50% shallower gradient over time (Figure S3B). Automated tracking of single-cell’s motility trajectories was performed for 14h in total. MCF7-EGFR^*mCitrine*^, as well as MCF10A cells migrated in a directional manner towards the EGF^647^ source (Figure 3A- and Figure S3C,D - left, green trajectory parts). This directed migration persisted for transient period of time after the gradient wash-out (Figure 3A- and Figure S3C,D - left, red trajectory parts, movie S3), indicating that cells maintain memory of the location of previously encountered source. After the memory phase, the cells transitioned to a migration pattern equivalent to that in the absence of a stimulus (Figure 3A right, Figures S3C,D middle). Uniform stimulation with 20ng/ml EGF^647^ did not induce directed migration in either of the cell lines, although the overall migration distance was increased (Brueggemann et al., 2021) (Figures S3C,D, right). Quantification of the directionality of single cells’ motion, that is defined as the displacement over travelled distance, showed that for MCF10A cells it was significantly higher during the gradient stimulation (5h) as compared to no- or uniform-stimulation case (Figure 3B). Moreover, the directionality estimated in the 9h time-frame after the gradient removal was greater than the one in continuous stimulus absence, corroborating that cells transiently maintain memory of the previous direction of migration.

**Figure 3.**
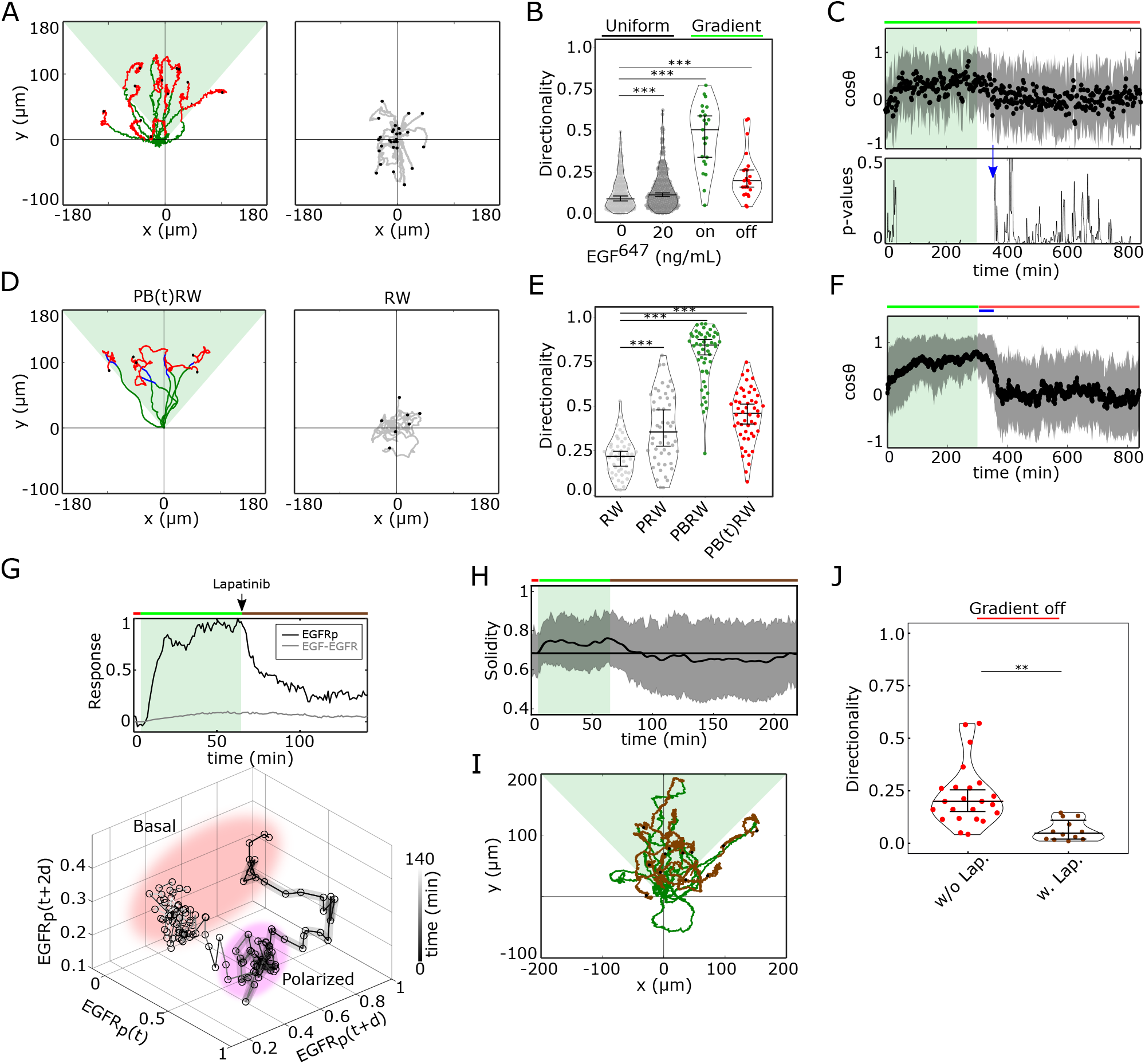
Cells display memory of recently encountered signals. **A**, Left: representative MCF10A single-cell trajectories. Green - 5h during and red line - 9h after dynamic EGF^647^ gradient (shaded). Exemplary cell in movie S3. Right: Same as in **A**, only 14h in continuous EGF^647^ absence. Black dots: end of tracks. **B**, Directionality (displacement/distance) in MCF10A single-cell migration during 14h absence (0ng/ml; n=249, N=3) or uniform 20ng/ml EGF^647^ stimulation (n=299, N=3); 5h dynamic EGF^647^ gradient (green) and 9h during wash-out (red; n=23, N=5). p-values: ***p ≤ 0.001, twosided Welch’s t-test. Error bars: median±95%C.I. **C**, Top: Projection of the cells’ relative turning angles (mean±sd; n=23, N=5) during (green shaded) and after 5h dynamic EGF^647^ gradient. Green/red lines: stimulus presence/absence. Bottom: Kolmogorov-Smirnov (KS) test p-values depicting end of memory in directional migration (arrow, *t* = 350*min*). KS-test estimated using 5 time points window. For **A**-C, data sets in Figures S3D, S4A-C. **D**, Representative *in silico* single-cell trajectories (Methods). Left: PB(t)RW: Persistent biased random walk, bias is a function of time (green/blue trajectory part - bias on). Right: RW: random walk. **E**,Corresponding directionality estimates from n=50, data in Figure S4D. PRW: persistent random walk. ***p-values: p ≤ 0.001, two-sided Welch’s t-test. Error bars: median±95%C.I. **F**, Same as in **C**, only from the synthetic PB(t)RW trajectories. **G**, Top: Exemplary profiles of *EGFR*_*p*_ (black) and *EGF* − *EGFR* (grey) in live MCF7-EGFR^*mCitrine*^ cell subjected to 1h EGF^647^ gradient (green shading), and 4h after wash-out with 1 µM Lapatinib. Mean±s.d. from n=9, N=2 in Figure S4H. Bottom: Corresponding reconstructed state-space trajectory with state-space trapping (colored) (Methods. movie S4). **H**, Average solidity in MCF7-EGFR^*mCitrine*^ cells subjected to experimental conditions as in **G**. Mean±s.d. from n=9, N=2 cells. **I**, MCF10A single-cell trajectories quantified 5h during (green) and 9h after (orange) dynamic EGF^647^ gradient (shading) wash-out with 3 µM Lapatinib. n=12, N=5. See also movie S5. **J**, Directionality in single-cell MCF10A migration after gradient wash-out with (brown, n=12, N=5) and without Lapatinib (red, n=23, N=5). p-values: ** p≤0.01, KS-test. Error bars: median±95%C.I. See also Figures S3 and S4.

This was also reflected in the projection of the cell’s relative turning angles (cos *θ*) estimated along the gradient direction (*π*) at each time point (Figure S4A), representing the angular alignment of the cells to the source direction. The cellular migration trajectories aligned with the source direction (cos *θ* approached 1) during, and maintained this temporally after gradient removal, before returning to a migration pattern characteristic for stimulus absence or during uniform stimulation (cos *θ* ≈ 0, Figure 3C top, Figure S4B). Calculating the similarity between the Kernel Density distribution Estimate (KDE) of the angular alignment distributions at each point in the gradient series with that in continuous stimulus absence, showed that the distributions approach each other only ∼ 50*min* after the gradient removal (Figure 3C, bottom). Additionally, the calculated similarity between the KDE distributions during the gradient (5h) and the 50*min* memory period further corroborated this finding (Figure S4C). The average memory phase in directional motility thus corresponds to the time-frame in which the memory in polarized EGFR^*mCitrine*^ phosphorylation and cell shape is maintained (Figures 2C-H), indicating that the metastable signaling state is translated to a stable prolonged migration response after gradient removal.

To investigate whether the motility patterns during the gradient and the memory phase have equivalent characteristics, we fitted the motility data using a modified Ornstein-Uhlenbeck process (Uhlenbeck and Ornstein, 1930; Svensson et al., 2017) and used the extracted migration parameters to generate synthetic single-cell trajectories (Methods). In absence of stimulus, the cellular motion resembled a random walk process (RW: Figure 3D right, Figures S4D,E middle), persistent random walk (PRW) was characteristic for the uniform stimulation case (Figure S4D, E right), whereas biased PRW described the migration in gradient presence (PBRW, Figure 3D- and Figure S4D, left, green trajectory part). Extending the bias duration during the interval of the experimentally observed memory phase (PB(t)RW) was necessary to reproduce the transient persistent motion after gradient removal (Figure 3D- and Figure S4D, left, blue trajectory part; Figures 3E,F; Figure S4F). Altogether, these results demonstrate that epithelial cells transiently maintain a memory of previous signal location and thereby display directed motility equivalent to that in the presence of a gradient, before reverting to a random walk migration pattern.

To corroborate the link between memory in polarized receptor activity and memory in directional migration, we quantified EGFR^*mCitrine*^ phosphorylation polarization in the MCF7-EGFR^*mCitrine*^, as well as directional migration of MCF10A cells, when cells were subjected to an ATP analog EGFR inhibitor Lapatinib (Bjorkelund et al., 2012) during gradient washout. The exemplary single-cell kymograph and EGFR^*mCitrine*^ phosphorylation temporal profile demonstrate that the phosphorylation response exponentially decays upon Lapatinib addition, resulting in a clear absence of transient memory in EGFR^*mCitrine*^ phosphorylation polarization (Figure 3G top, Figures S4G,H). This is further reflected in the reconstructed state-space trajectory that smoothly transits from the polarized to the basal activity state, without the transient state-space trapping that was characteristic for the memory state emerging from the dynamical ”ghost” (compare Figure 3G bottom to 2F, movie S4). The absence of memory in EGFR^*mCitrine*^ phosphorylation was also reflected in absence of transient memory in morphological changes after stimulus removal (Figure 3H). In the MCF10A migration assay, cells directly switched to RW migration pattern upon gradient wash-out with Lapatinib, as shown through the directionality quantification after gradient removal (Figures 3I,J; movie S5). Equivalent single-cell motility trajectories could be mimicked with the PB(t)RW simulation, where the bias duration corresponded to the duration of the gradient (Figures S4I,E). This shows that the transient memory arising from a metastable ”ghost” signaling state is a core dynamical feature underlying transient memory in directional motility, and cannot be explained with slow relaxation kinetics in receptor dephosphorylation.

### 4 Molecular working memory enables cells to navigate in dynamic chemoattractant fields

To test whether the identified memory enables cellular navigation in environments where signals are disrupted but also change over time and space, we subjected cells in the simulations and experiments to a changing growth factor field. The field was generated by a sequence of signals, starting with a dynamic gradient whose steepness changed over time, that was temporary disrupted for a time interval shorter than the interval of memory in cell polarization, followed by a second static gradient in the same direction, that after an equivalent disruption period was followed by a third dynamic gradient in the opposite direction (Figure 4A). The *in silico* migration simulations showed that the cell can sense the initial dynamic gradient and polarizes in the direction of maximal attractant concentration, resulting in directed migration (Figure 4B, Figure S5A, movie S6). The simulations also predicted that the memory of the previously encountered signal localization enables maintaining robust directional migration even when the signal was disrupted, while still remaining sensitive to the newly emerging signal from the opposite direction. In the simulation, the *in silico* cell rapidly adapted the orientation when encountering the third signal with opposite localization, demonstrating that the proposed mechanism can also account for prioritizing newly encountered signals. Such a dynamic memory which enables information of previous signals to be temporally maintained while retaining responsiveness to upcoming signals and thereby manipulate the stored information, in neuronal networks is described as a working memory (Atkinson and Shiffrin, 1968). On the other hand, the simulations also showed that the long-term memory resulting from organization in the stable inhomogeneous steady state regime, hindered cellular adaptation to a changing gradient field. The initial dynamic gradient shifted the system to the stable polarization steady state where it was maintained in a long-term, such that sensitivity to upcoming signals from the same direction was hindered. Even more, in the simulations, the cell could not resolve the conflicting information from a subsequent gradient from the opposite direction as the signals induced high receptor activity on the opposed cell sides, resulting in ending the migration (Figures S5B,C, movie S7).

**Figure 4.**
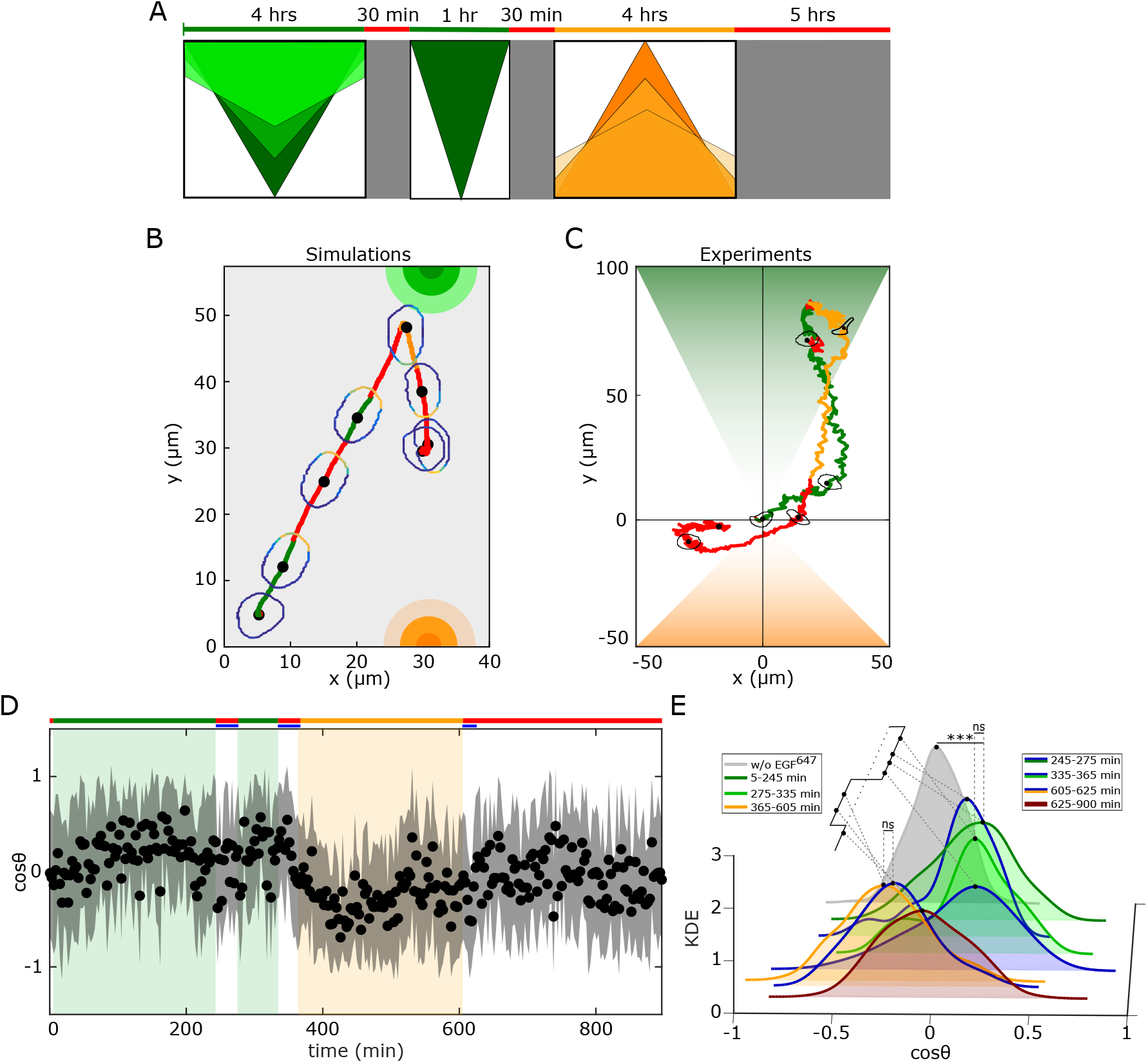
History-dependent single-cell migration in changing chemoattractant field. **A**, Scheme of dynamic spatial-temporal growth factor field implemented in the simulations and experiments. Green(orange)/red: gradient presence/absence. **B**, *In silico* cellular response to the sequence of gradients as depicted in **A**, showing changes in EGFR activity, cellular morphology and respective motility trajectory over time. Trajectory color coding corresponding to that in (**A**), cell contour color coding with respective *E*_*p*_ values as in Figure 1E. Cell size is magnified for better visibility. See also movie S6. **C**, Representative MCF10A single-cell trajectory and cellular morphologies at distinct time-points, when subjected to dynamic EGF^647^ gradient field as in **A**. Trajectory color coding corresponding to that in **A**. See also movie S8. **D**, Projection of cells’ relative turning angles (cos *θ*) depicting their orientation towards the respective localized signals. Mean±s.d. from n=12, N=5 is shown. Data in Figure S5E. **E**, Corresponding kernel density estimates (intervals and color coding in legend). p-values:, ***p ≤ 0.001, ns: not significant, KS-test. See also Figure S5.

We next tested these predictions experimentally, by establishing an equivalent dynamic EGF^647^ spatial-temporal field in a controlled manner in the microfluidic chamber, and quantified the migratory profile of MCF10A cells (Figure S5D). The MCF10A cells sensed the initial dynamic gradient field and migrated in the direction of the largest chemoattractant concentration, maintaining the directionality even when the signal was temporary disrupted. Despite the memory in cell polarization, cells remained responsive and adapted the duration of directional migration when presented with a second static gradient from the same direction, and subsequently prioritized the third, newly encountered signal with opposed orientation (exemplary trajectory in Figure 4C, movie S8, Figure S5E). The temporal memory in directional migration as well as the continuous adaptation of MCF10A cells to novel cues was also reflected in the projection of the cell’s relative turning angles (Figure 4D), whereas the respective KDE distributions derived from the subsequent time-intervals of gradient presence/absence corroborate that cells maintain the same migration characteristics within the memory intervals as during the gradient phase (Figure 4E). These results demonstrate that cells navigate in changing gradient fields by utilizing a molecular mechanism of working memory that is an intrinsic feature of receptor tyrosine kinase networks.

## Discussion

Our data establishes that mammalian cells use a mechanism of working memory to navigate in complex environments where the chemical signals are disrupted or vary over time and space. Even though persistent migration of eukaryotic cells in absence of signals was previously observed (Skoge et al., 2014; Albrecht and Petty, 1998; Prentice-Mott et al., 2016), its underlying mechanism and the implications for navigation in changing environments have not been elucidated. The mechanism of transient memory we report here is realized on a molecular level, by storing information about direction of previously encountered signals through maintaining a prolonged polarized phosphorylation state of receptor tyrosine kinases. Dynamically, the prolonged polarized state emerges for organization of receptor networks at criticality, where a slow-escaping remnant from the attractor state or a dynamical ”ghost” is generated. The ”ghost” maintains the system away from steady state, suggesting that in migrating cells, the information about previously encountered signals can be encoded in the transient state-space trajectories rather than the steady-states of the protein interaction network. Our simulations and migration experiments show that this encoding via transient states is necessary to ensure the ability of cells to adapt to changes in the external environment, while maintaining memory of previous signals. Our work furthermore suggest that this general mechanism of a system poised at criticality can explain a wide range of biologically relevant scenarios, from the integration of temporally and spatially varying signals, to how extracellular information is transformed into guidance cues for memory-directed migration. Memory-guided navigation is advantageous when migration must be realized over long and complex trajectories through dense tissues where the chemical cues are disrupted or only locally organized.

For neuronal networks, short-term memory is a main requirement to integrate temporal dependencies from changing signals (Hochreiter and Schmidhuber, 1997; Maass et al., 2000). We have demonstrated here that the transient memory in cell polarization and therefore the capabilities of cells to navigate in a complex environment are an emergent feature of receptor networks organized at criticality, and cannot be explained using computations with stable states. It would be of interest to study whether receptor networks are self-organized at criticality, or these features arose through evolution as a means for optimizing the computational capabilities of cells. The identification of a molecular working memory also opens avenues of research in the single-cell migration and tissue homeostasis to study whether cells can integrate and interpret even sub-threshold environmental signals, leading to release of cells from a tissue and long-distance single cell migration, as during cancer metastasis.

## Acknowledgments

The authors thank Angel Stanoev for initial analysis of the EGFR polarity model, critical discussion during the project, as well as valuable comments on the manuscript, Manish Yadav and Monika Scholz for critical reading of the manuscript and valuable suggestions, and Frédéric Paquin-Lefebvre for valuable suggestions on the realization of the reaction-diffusion simulations. All of the experiments were carried in the lab of Philippe Bastiaens, and we are particularly grateful for the opportunity to be part of that engaging and critical community where we could learn and develop this project. We especially thank P. Bastiaens for numerous critical discussion and suggestionsthat were crucial throughout the project, as well as for detailed comments that significantly helped us to improve the manuscript.

## Data and code availability

The data can be obtained from the corresponding author upon reasonable request. The codes will be made publicly available upon acceptance.

## Funding

The project was funded by the Max Planck Society, partially through the Lise Meitner Excellence Program.

## Author contributions

A.K. conceptualized and supervised the project; A.P.N. and A.K. developed the theoretical description; A.P.N. performed the analytical and numerical analysis; A.D. performed most of the experiments with help of A.P.N.; A.D., A.P.N. and R.L. analyzed the data. A.K. wrote the manuscript with help of A.D., A.P.N. and R.L.

## Competing Interests statement

The authors declare no competing interests.

## 5 Materials and Methods

### 5.1 Cell Culture

MCF7 cells (sex: female, ECACC, Cat. No. 86012803) were grown at 37°C and 5% *CO*_2_ in Dulbecco’s Eagle’s medium (DMEM) (PAN-Biotech, Germany), supplemented with 10% inactivated Fetal Calf Serum (FCS) (Sigma-Aldrich), 100 ng ml^−1^ L-Glutamine, 0.5 mg ml^−1^ non-essential amino acids, 100 µg ml^−1^ penicillin and 100 µg ml^−1^ streptomycin (PAN-Biotech, Germany). Serum starvation was performed by culturing the cells in DMEM supplemented with 0.5% FCS, 100 µg ml^−1^ penicillin and 100 µg ml^−1^ streptomycin (PAN-Biotech, Germany). MCF10A cells (sex: female, ATCC-CRL 10317) were grown at 37°C and 5% *CO*_2_ in Mammary Epithelial Cell Growth Basal medium (MEBM from Lonza Pharma & Biotech), supplemented with 5% Horse Serum (HS) (Invitrogen), 20 ng mL^−1^ EGF (Sigma-Aldrich), 0.5 mg mL^−1^ hydrocortisone (Sigma-Aldrich), 100 ng ml^−1^ cholera toxin (Sigma-Aldrich), 10 µg mL^−1^ insulin (Sigma-Aldrich), 100 µg mL^−1^ penicillin and 100 µg mL^−1^ streptomycin. Serum starvation was performed by culturing the cells in the DMEM supplemented with 0.5% HS, 0.5 mg mL^−1^ hydrocortisone (Sigma-Aldrich), 100 ng ml^−1^, cholera toxin (Sigma-Aldrich) 100 µg mL^−1^ penicillin and 100 µg mL^−1^ streptomycin. MCF7 and MCF10A cells were authenticated by Short Tandem Repeat (STR) analysis and did not contain DNA sequences from mouse, rat and hamster (Leibniz-Institut DSMZ). Cells were regularly tested for mycoplasma contamination using MycoAlert Mycoplasma detection kit (Lonza).

### 5.2 Transfection and cell seeding

For EGFR^*mCitrine*^ polarization experiments, 2.5 × 10^5^ MCF7 cells were seeded per well in a 6-well Lab-Tek chamber (Nunc) until 80% confluence was reached. After 9-10 h of seeding, transient transfection was performed with a total of 1 µg of plasmids (*EGFR*^*mCitrine*^, *PTB*^*mCherry*^ and *cCbl*^*BF P*^ at ratio 4:3:4 by mass) using FUGENE6 (Roche Diagnostics) transfection reagent and Opti-MEM (Gibco - Thermo Fisher Scientific) according to manufacturer’s procedure. All plasmids were generously provided by Prof. P. Bastiaens, MPI of Molecular Physiology, Dortmund. Cells were incubated for 7-8 h to allow the expression of the transfected proteins prior to experiments. To detach the cells, the growth media was discarded and cells were washed once with DPBS (PAN Biotech) before adding 100 µL Accutase (Sigma-Aldrich). After 10 min incubation period at 37°C and 5 % CO2, fresh growth media was added, and the cell density and viability was measured using cell counter (Vi-CELL XR Cell Viability Analyzer System). After spinning down, the cells were diluted to 10 × 10^6^ cells/ml. The M04-G02 microfluidic gradient plates (Merck Chemicals) were primed for usage by flowing cell culture growth media through the cell chamber for 5 min and cells were subsequently seeded according to manufacturer’s instructions.

For migration experiments with uniform *EGF* ^647^ stimulation, 6-well Lab-Tek plates were coated with Collagen (Sigma-Aldrich) in 0.1 M Acetic acid (Sigma-Aldrich) for MCF7 (100 µg cm^−2^), and Fibronectin (Sigma-Aldrich) in Phosphate-Buffered Saline (DPBS) (PAN-Biotech) for MCF10A cells (2 µg mL^−1^), and stored in incubator at 37°C overnight for evaporation. Excessive media was removed and the wells were washed with DPBS before seeding cells. MCF7 cells were seeded and transfected as described above. In the case of MCF10A cells, 1 × 10^5^ cells per well were used for seeding. For migration experiments with gradient EGF^647^ stimulation, MCF7 cells were transferred to the coated M04-G02 microfluidic gradient plates as described above. Before seeding, MCF10A cells were detached from 6 well Lab-Teks by discarding the growth media and washing once with DPBS (PAN Biotech) before adding 100 µL Accutase (Sigma-Aldrich). After 20 − 30*min* incubation period at 37°C and 5 % CO2, fresh cell growth media was added, and the cell density and viability were measured using a cell counter (Vi-CELL XR Cell Viability Analyzer System). After spinning down, the cells were diluted to 2 × 10^6^ cells/ml, and subsequently seeded in the microfluidic plates according to manufacturer’s instructions.

### 5.3 Reagents

For gradient quantification, Fluorescein (Sigma Aldrich) was dissolved in Dulbecco’s modified Eagle’s medium (with 25 mM HEPES, without Phenol Red) (PAN Biotech). Imaging media: DMEM without Phenol Red was mixed with 25 mM HEPES. For nuclear staining, 20 mM Hoechst 33342 (Thermo Fisher Scientific) was mixed with DPBS and diluted to 2 µM working concentration. EGFR inhibitor Lapatinib (Cayman Chemical, Ann Arbor, MI) was solubilized in DMSO (Thermo Fisher Scientific) to a stock concentration of 5 mM and stored at -20°C.

### 5.4 Confocal and wide-field microscopy

Confocal images were recorded using a Leica TCS SP8i confocal microscope (Leica Microsystems) with an environment-controlled chamber (Life Imaging Services) maintained at 37°C and HC PL APO 63x/1.2 N.A / motCORR CS2 water objective (Leica Microsystems) or a HC PL FLUOTAR 10x/0.3 N.A. dry objective (Leica Microsystems). mCitrine, mCherry and Alexa647 were excited with a 470 nm-670 nm pulsed white light laser (Kit WLL2, NKT Photonics) at 514 nm, 561 nm and 633 nm, respectively. BFP and Hoechst 33342 (Thermo Fisher Scientific) were excited with a 405 nm diode laser. The detection of fluorescence emission was restricted with an Acousto-Optical Beam Splitter (AOBS): BFP (425 nm-448 nm), Hoechst 33342 (425 nm-500 nm), mCitrine (525 nm-551 nm), mCherry (580 nm-620 nm) and Alexa647 (655 nm-720 nm). Transmission images were recorded at a 150-200% gain. To suppress laser reflection, Notch filter 488/561/633 was used whenever applicable. When using the dry objective for migration experiments, the pinhole was set to 3.14 airy units and 12-bit images of 512×512 pixels were acquired in frame sequential mode with 1x frame averaging. When using the water objective for polarization experiments, the pinhole was fixed (1.7 airy units) for all channels. The Leica Application Suite X (LAS X) software was used.

Wide field images were acquired using an Olympus IX81 inverted microscope (Olympus Life Science) equipped with a MT20 illumination system and a temperature controlled *CO*_2_ incubation chamber at 37°C and 5% *CO*_2_. Fluorescence and transmission images were collected via a 10x/0.16 NA air objective and an Orca CCD camera (Hamamatsu Photonics). Hoechst 33342 fluorescence emission was detected between 420 nm-460 nm via DAPI filter, mCitrine fluorescence emission between 495 nm-540 nm via YFP filter and Alexa647 fluorescence emission between 705 nm-745 nm via Cy5 filter. The xCellence (Olympus) software was used.

### 5.5 Gradient establishment for polarization and migration experiments

The CellAsic Onix Microfluidic Platform (EMD Millipore) was used for gradient cell migration and EGFR^*mCitrine*^ phosphorylation polarization experiments. For EGFR^*mCitrine*^ phosphorylation polarization experiments, 1 h gradient stimulation was established using CellASIC ONIX2 software as follows. (i) Pre-stimulus: Imaging media was flowed from well groups 3 and 4 (CellAsic Onix Manual - www.merckmillipore.com/) at low pressure (2.5 kPa) for 5 min. (ii) Gradient establishment: After closing well group 3, pre-loaded EGF^647^ (10 ng mL^−1^) was flowed through well group 2 and imaging media from well group 4 at high pressure (15 kPa) for 15 min (iii) Gradient maintenance: The pressure was reduced to 10 kPa for 45 min. (iv) Washout: After closing well groups 2 and 4, imaging media was flowed from well groups 3 and 5 at high pressure (15 kPa) for 15 min and maintained at low pressure (7 kPa) for 165 min. For single gradient migration experiments, this protocol was modified as follows: in step (iii), gradient maintenance was done for 285 min. In step (iv), maintenance was at low pressure for 585 min. 30 ng mL^−1^ EGF^647^ was used. For polarization experiments with inhibitor, the same protocol as for polarization experiments was used, except well group 3 and 5 were filled with 1 µM Lapatinib solution and in step (i) well group 3 was kept closed. For single cell gradient migration experiment with inhibitor, 3 µM Lapatinib was used.

For migration experiments under subsequent gradient stimuli / gradient quantification, the following changes in the steps were used : (ii) well group 2 with 30 ng mL^−1^ EGF^647^/ 2.5 µM Fluorescein was used. (iii) The gradient maintenance was done for 225 min. (iv) Washout: imaging media was flowed from well groups 3 and 4 at high pressure (15 kPa) for 15 min and maintained at low pressure (7 kPa) for 15 min. (v) Second gradient establishment: After closing well group 3, EGF^647^(30 ng mL^−1^) / 2.5 µM Fluorescein was flowed from well group 2 and imaging media from well group 4 at high pressure (15 kPa) for 15 min. (vi) The second gradient thus formed was maintained by reducing the pressure to 10 kPa for 45 min. (vii) Washout: imaging media was flowed from well groups 3 and 4 at high pressure (15 kPa) for 15 min and maintained at low pressure (7 kPa) for 15 min. (viii) Third gradient establishment: After closing well group 4, EGF^647^ (30 ng mL^−1^) / 2.5 µM Fluorescein was flowed from well group 5 and imaging media from well group 3 at high pressure (15 kPa) for 15 min. (ix) The third reversed gradient was maintained by reducing the pressure to 10 kPa for 225 min. (x) Washout: imaging media was flowed from well groups 3 and 4 at high pressure (15 kPa) for 15 min and maintained at low pressure (7 kPa) for 285 min.

### 5.6 Imaging *EGFR*^*mCitrine*^ phosphorylation polarization and single cell migration

Transfected MCF7-EGFR^*mCitrine*^ cells transferred to M04G-02 gradient plates as described above were incubated for at least 3 h, followed by serum starvation for at least 6 h before imaging. Existing cell media was substituted right before imaging with imaging media. Confocal imaging for multiple positions at 1 min time interval using adaptive auto-focus system and the water objective was performed concurrently during the duration of the experiment using the Leica TCS SP8i.

For migration experiments under uniform EGF^647^ stimulation, confocal laser scanning microscopy / transmission imaging of live MCF7-EGFR^*mCitrine*^ / MCF10A cells was done on a Leica TCS SP8i or Olympus IX81 for multiple positions at 3 min and 2 min time interval respectively, using the 10x dry objective for 14 hours.

### 5.7 EGF^647^ / Fluorescein gradient quantification

hEGF^647^ was generated in the lab of Prof. P. Bastiaens, MPI of molecular Physiology, Dortmund, using the His-CBD-Intein-(Cys)-hEGF-(Cys) plasmid (Sonntag et al., 2014), kindly provided by Prof. Luc Brunsveld, University of Technology, Eindhoven. Human EGF was purified from E. coli BL21 (DE3), N-terminally labeled with Alexa647-maleimide as described previously (Sonntag et al., 2014) and stored in PBS at -20°C. To quantify the spatial extent of the EGF^647^ / Fluorescein gradient, gradients were generated following the protocol described in sub-section 5.5 in plates without cells or matrix coating. Confocal images of Alexa647 / GFP channel were acquired at 1 min interval. A rectangular region of interest (including the perfusion channels and the culture chamber) was used to obtain an averaged pixel intensity profile using FIJI at each time point. This spatial profile was averaged across multiple experiments and then scaled with the mean intensity value in the perfusion channel, which corresponds to the applied EGF^647^ / Fluorescein concentration.

### 5.8 Quantifying EGFR^*mCitrine*^ phosphorylation in single cells

To quantify plasma membrane EGFR^*mCitrine*^ phosphorylation in live MCF7-EGFR^*mCitrine*^ cells, single cell masks were obtained from the EGFR^*mCitrine*^ channel at each time-point using FIJI (https://imagej.net/Fiji). All pixels within the obtained boundary were radially divided into 2 segments of equal areas (Stanoev et al., 2018), and the outer segment was taken to represent the plasma membrane. For the kymograph analysis, at each time point, the plasma membrane segment was divided into 4 quadrants in anti-clockwise direction, and each was divided into 5 spatial bins (Figure 2A). The fraction of phosphorylated EGFR^*mCitrine*^ in each bin, *i* was estimated as:

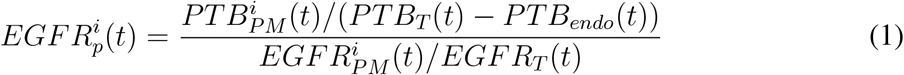

where 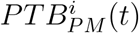 and 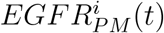 are respectively the PTB^*mCherry*^ and *EGFR*^*mCitrine*^ fluorescence at *i*^*th*^ plasma membrane bin, *PTB*_*T*_ (*t*) and *EGFR*_*T*_ (*t*) - respective total fluorescence in the whole cell, *PTB*_*endo*_(*t*) – the PTB^*mCherry*^ fluorescence on vesicular structures in the cytoplasm. Endosomal structures were identified from the cytosol by intensity thresholding (1.5 s.d. percentile) and PTB^*mCherry*^ fluorescence from these structures was subtracted from the *PTB*_*T*_ (*t*), to correct for the PTB^*mCherry*^ fraction bound to the phosphorylated EGFR^*mCitrine*^ on endosomes.

Temporal profile of the fraction of phosphorylated EGFR^*mCitrine*^ on the plasma membrane was obtained using:

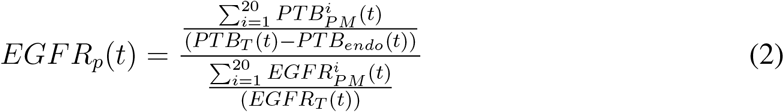

and then normalized as:

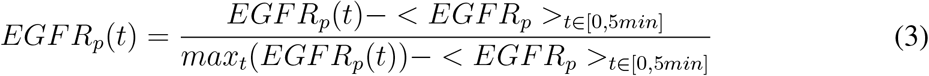

with <> being the temporal average in the pre-stimulation interval *t* ∈ [0, 5*min*]. The fraction of liganded receptor was calculated using:

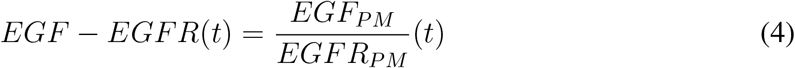

To classify single cells into non-activated, activated (polarized EGFR^*mCitrine*^ phosphorylation) and pre-activated (uniformly distributed EGFR^*mCitrine*^ phosphorylation) upon gradient EGF^647^ stimulation (Figure S2A, B), the following method was applied. To identify preactivated cells, a Gaussian Mixture Model (GMM) was fitted to the histogram of 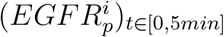 values from all the analysed cells, and the intersection point between the two normal distributions was identified. If more than 30% of the 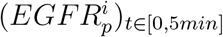 pixel intensity values for any cell lie above the intersection point, the cell is classified as pre-activated. To distinguish between the non-activated and activated cells in the remaining population, average EGFR^*mCitrine*^ phosphorylation value (*EGFR*_*p*_) per cell was estimated during the pre-stimulation (*t* ∈ [0, 5*min*]) and the stimulation period (*t* ∈ [5*min*, 65*min*]) (*< EGFR*_*p*_ *>*_*t*∈[0,65]_) from the temporal EGFR^*mCitrine*^ phosphorylation profiles. Histogram of the respective *EGFR*_*p*_ values was again fitted with a GMM model. All cells with an average *< EGFR*_*p*_ *>*_*t*∈[0,65]_ value lying below the intersection point were considered to be non-activated, whereas those above - activated.

The average of the spatial projection of the fraction of phosphorylated EGFR^*mCitrine*^ from single-cell kymographs (Figure S2C) was generated from the 20 (from total of 21 cells) that were polarized in the direction of the EGF^647^ gradient. For each cell, a temporal average of *EGFR*_*p*_ per bin was calculated for the duration of the gradient (*t* ∈ [5*min*, 65*min*]) and the bin with the maximal *EGFR*_*p*_ value was translated to *π*. The profiles were then smoothened using a rolling average with a window of 7 bins. The resulting profiles were then averaged over all cells and mean±s.d. is shown.

### 5.9 Estimating memory duration in EGFR^*mCitrine*^ phosphorylation polarization

The duration of memory in EGFR^*mCitrine*^ phosphorylation polarization in single cells was estimated from the temporal profile of the fraction of plasma membrane area with high EGFR^*mCitrine*^ phosphorylation during and after gradient removal (Figures 2D,E). For this, the single-cell kymographs were normalized to a maximal value of 1 using

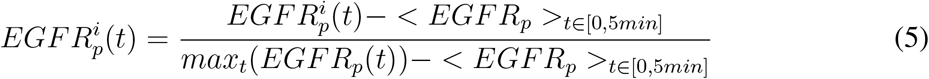

yielding the value of phosphorylated EGFR^*mCitrine*^ per bin *i* per time point *t*. Using the mean of *EGFR*_*p*_ + *s*.*d*. over the whole experiment duration as a threshold, all 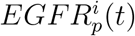 lying above the threshold were taken to constitute the area of polarized EGFR^*mCitrine*^ phosphorylation. To account for different bin sizes, at each timepoint, the area of all bins with *EGFR*_*p*_ above the threshold was summed and divided by the respective total cell area, yielding the temporal evolution of the fraction of polarized cell area (FPA) (Figure 2D). The end of the memory duration per cell was identified as the time point at which *FPA*_*per*−*cell*_ *<* (*FPA*_*average*_ − *s*.*d*.) in 3 consecutive time points.

### 5.10 Quantifying morphological changes in response to EGF^647^ in experiments and simulations

Morphological changes of polarized cells were quantified using the solidity (Figure 2H) of each cell at each time point and the directed protrusive area towards and away from the gradient (Figure 1 G,H; 2G). The solidity *σ* is the ratio between the cell’s area *A*_*cell*_ and the area of the convex hull 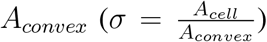. The memory duration in cell morphology was calculated from the single-cell solidity profiles, and corresponds to the time-point at which the solidity is below mean-s.d. estimated during gradient presence. The directed cell protrusion area was estimated by comparing single cell masks at two consecutive time points. To reduce noise effects, the masks were first subjected to a 2D Gaussian filtering using the *filters*.*gaussian* function from the *scipy* python package. Protrusions were considered if the area change was greater than 10 pixels or 1.2*µm*^2^ per time point. The front and the back of the cells were determined by identifying an axis that runs perpendicular to the gradient and through the cell nucleus of the initial time point. The directed cell protrusion area was then obtained using 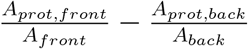. The final profiles of directed protrusive area were smoothed using 1D Gaussian filtering with the *filters*.*gaussian filter*1*d* function from the *scipy* python package. For the equivalent quantification from the simulations, the same procedures were applied without an area threshold. The memory duration was estimated as the time point at which the directed protrusive area crosses zero after the gradient removal.

### 5.11 Quantification of single-cell migration and duration of memory in migration

Single cell migration trajectories were extracted using Trackmate (Tinevez et al., 2017) in Fiji (Schindelin et al., 2012) using Hoechst 33342 / transmission channel. From the positional information (x and y coordinates) of individual cell tracks, quantities such as Motility, Directionality and cos *θ* were extracted using custom made Python code (Python Software Foundation, versions 3.7.3, https://www.python.org/). Directionality was calculated as displacement over total distance and statistical significance was tested using two-sided Welch’s t-test. To quantify the memory duration in directed single-cell migration, the Kernel Density Estimate (KDE) from cos *θ* quantification in the continuous absence of EGF^647^ (uniform case, between 250 min-300 min) was compared with windowed KDE (5 time points moving window) from the gradient migration profile, using two sided Kolmogorov-Smirnov test.

To quantify the motility patterns of MCF10A cells in absence, uniform or gradient *EGF* ^647^ stimulation, we fitted the experimentally obtained single cell migration trajectories using modified Ornstein-Uhlenbeck process (mOU) (Uhlenbeck and Ornstein, 1930) that is defined by the Langevin equation for the velocity vector *ν*:

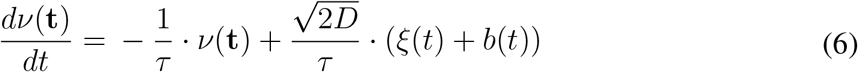

where *ξ*(*t*) represents a white noise component, D is a diffusion coefficient characteristic of a Brownian motion, *τ* is the persistence time and *b*(*t*) models the contribution of the time-dependent bias. The experimental data was fitted to obtain values of D and *τ*. In order to estimate D, Mean Square Displacement (MSD) was calculated from the single cell tracks using 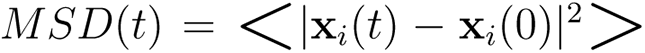, where **x**_*i*_(*t*) is the tracked position of *i*-th cell in the 2D plane, *<>* is the average across all single cell tracks, and |.| is the Euclidean distance (D. et al., 2005). To estimate D, the obtained MSD profile was fitted with a linear function (= 4*Dt*). Goodness of Fit for the different experimental conditions: 0ng/ml EGF^647^, *R*^2^ = 0.975; for uniform 20ng/ml EGF^647^ stimulation, *R*^2^ = 0.995. In order to estimate *τ*, Velocity Auto-Correlation Function 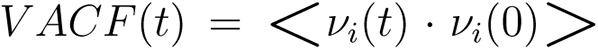, where *ν*_*i*_(*t*) is the measured velocity of *i*-th cell at time t, was fitted with a mono exponential function 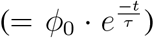. Goodness of Fit : for 0ng/ml EGF^647^ case - Standard Error Of Estimate *SEOE* = 0.0261; for uniform 20ng/ml EGF^647^ stimulation case, *SEOE* = 0.0570. Fitted values: for 0ng/ml EGF^647^ case, *τ* = 11.105, *D* = 0.425; for uniform 20ng/ml EGF^647^ stimulation case, *τ* = 38.143, *D* = 2.207; bias *b*(*t*) = 0.134.

### 5.12 Reconstructing state-space trajectories from temporal EGFR^*mCitrine*^ phosphorylation profiles

The state-space reconstruction in Figures 2F and 3G was performed using the method of timedelay. For a time series of a scalar variable, a vector *x*(*t*_*i*_), *i* = 1,..*N* in state-space in time *t*_*i*_ can be constructed as following

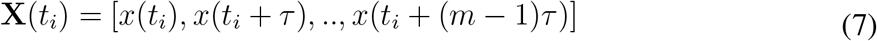

where *i* = 1 to *N* −(*m* − 1)*τ, τ* is the embedding delay, *m* - is a dimension of reconstructed space (embedding dimension). Following the embedding theorems by Takens (Takens, 1980) and Sauer et al. (Sauer et al., 1991), if the sequence *X*(*t*_*i*_) consists of scalar measurements of the state of a dynamical system, then under certain genericity assumptions, the time delay embedding provides a one-to-one image of the original set, provided *m* is large enough. The embedding delay was identified using the *timeLag* function (based on autocorrelation), the embedding dimension using the *estimateEmbeddingDims* function (based on the nearest-neighbours method), and the state-space reconstruction using the *buildTakens* function, all from the *nonlinearTseries* package in R (https://cran.r-project.org/web/packages/nonlinearTseries/index.html). Before state-space reconstructions, time series were smoothened using the *Savitzky-Golay* filter function in Python. For Figure 2F, *τ* = 26, *d*_*e*_ = 3; for Figure 3G, *τ* = 50, *d*_*e*_ = 3.

## Notes

### Competing Interest Statement

The authors have declared no competing interest.

