## Supplementary information for "Cells use molecular working memory to navigate in changing chemoattractant fields"

### Supporting Information

#### 0.1 Theoretical consideration of the navigation mechanism in a generalized reaction-diffusion signaling model

We consider a generalized form of a (mass-conserved) reaction-diffusion (RD) model of an $M$ ($\mathbf{U} \in \mathbf{R}^M$) component system in $N$ ($\mathbf{x} \in \mathbf{R}^N$) dimensional space

$$\frac{\partial \mathbf{U}(\mathbf{x}, t)}{\partial t} = \mathbf{F}(\mathbf{U}(\mathbf{x}, t)) + \mathbf{D} \cdot \nabla^2 \mathbf{U}(\mathbf{x}, t) \quad (1)$$

where $\mathbf{F} \in \mathbf{R}^M$ is the reaction term, $\mathbf{D}$ is a $M \times M$ diagonal matrix of diffusion constants $D_j, j = 1, ..., M$, and $\nabla^2$ is the Laplacian operator. Standard analysis of such models relies on linear stability analysis to find the conditions for a Turing-type instability (Turing, 1952), such that the symmetric steady state becomes unstable and an asymmetric polarized state is stabilized. By its nature, the linear stability analysis makes no prediction about the transition process itself, and thereby the type of bifurcation that underlies it. To provide quantitative description of the symmetry breaking transition in reaction-diffusion models, local perturbation analysis can be applied (Holmes et al., 2015). However, this analysis is mainly restricted to models characterized with large diffusion discrepancy between the signaling components. The conditions for a pitchfork bifurcation ($PB$)-induced transition in a generic RD model therefore have to be formally defined. Let $\mathbf{U_s} = (u_{is})$ for $i = 1, .., M$, be the stable homogeneous symmetric steady state of the RD system. Consider a linear perturbation of the form

$$\mathbf{U}(\mathbf{x}, t) = \mathbf{U_s} + \delta\mathbf{U}(\mathbf{x})e^{(\lambda t)}, \quad \delta\mathbf{U}(\mathbf{x}) \in \mathbf{R}^M \tag{2}$$

where $\delta\mathbf{U}(\mathbf{x})$ is the spatial and $e^{(\lambda t)}$ is the temporal part of the perturbation. Substituting Eq.(2) in Eq.(1) yields a linearized eigenvalue equation whose solution can be determined by solving the characteristic equation, $F_\lambda = det(\lambda I_{M\times M} - J_{M\times M}) = 0$. $J$ is the Jacobian matrix of the system defined by $J_{ij} = \frac{\partial \mathbf{F}_i(\mathbf{U}(x,t))}{\partial U_j}, i = 1, ...., M, j = 1, ....., M$.

The system exhibits a $PB$ if, an odd eigenfunction $\delta\mathbf{U}(\mathbf{x})$ such that $\delta\mathbf{U}(-\mathbf{x}) = -\delta\mathbf{U}(\mathbf{x})$, taken in the limit $\lambda \to 0$, fulfills the following condition (Paquin-Lefebvre et al., 2020):

$$\lim_{\lambda\to 0} F_\lambda = det(J) = 0. \tag{3}$$

When this conditions is satisfied, the symmetric, homogeneous steady state of the system undergoes a pitchfork bifurcation and an inhomogeneous steady state (IHSS) with two branches of asymmetric steady states emerges. In terms of polarization, these branches correspond to front-back-polarized states, where the orientation depends on the direction of the external signal (Figure 1A).

To identify whether the PB is of sub-critical type, and thereby identify the presence of a $SN_{PB}$, a weakly nonlinear analysis of Eq.(1) must be performed to obtain description of the amplitude dynamics of the inhomogeneous state. This can be achieved using an approximate analytical description of the perturbation dynamics based on the Galerkin method (Becherer et al., 2009; Rubinstein et al., 2012; Bozzini et al., 2015). For simplicity, we outline the steps for a one-dimensional system ($N = 1$). As we are interested in the description of a structure of finite spatial size (i.e. finite wavelength $k$), the final solution of the PDE is expanded around the fastest growing mode, $k_m$ into a superposition of spatially periodic waves. That means that $u(x, t) \in \mathbf{U}$ can be written as:

$$u(x, t) \approx \sum_{n=-\infty}^{+\infty} (u_n(t)e^{nik_m x} + u_n^*(t)e^{-nik_m x}) \quad (4)$$

where $u_n(t)$ is the complex amplitude of the $n^{th}$ harmonics. Let the amplitude corresponding to the leading harmonics ($n = 1$) is $\phi(t)$. After assuming that the amplitude of every other harmonics can be written as a power series of $\phi(t)$, substituting Eq.(4) into Eq.(1) allows to write an equation that describes the evolution of $\phi(t)$. In the case when the resulting equation is of Stuart-Landau type:

$$\frac{d\phi}{dt} = c_1\phi + c_2\phi^3 - c_3\phi^5 \quad (5)$$

with $c_1, c_2, c_3 > 0$, this corresponds to the normal form of a sub-critical pitchfork bifurcation (Strogatz, 2018). Together with the condition given by Eq.(3), the existence of a sub-critical PB for the full system (Eq.(1)) is guaranteed. A numerical or analytical analysis of Eq.(5) enables the identification of the position of the $SN_{PB}$.

#### 0.2 Modeling EGFR phosphorylation polarization dynamics

The dynamics of the experimentally identified spatially-distributed EGFR sensing network (Figure 1B, Figure S1A) is described using the following one-dimensional system of partial differential equations (PDEs):

$$
\begin{aligned}
\frac{\partial [E_p]}{\partial t} &= f_1([E_p], [E-E_p], [RG_a], [N2_a], [EGF_t]) + D_{E_p}\frac{\partial^2 [E_p]}{\partial x^2}\\
\frac{\partial [E-E_p]}{\partial t} &= f_2([E_p], [E-E_p], [EGF_t]) + D_{E-E_p}\frac{\partial^2 [E-E_p]}{\partial x^2}\\
\frac{\partial [RG_a]}{\partial t} &= f_3([E_p], [E-E_p], [RG_a]) + D_{RG_a}\frac{\partial^2 [RG_a]}{\partial x^2}\\
\frac{\partial [N2_a]}{\partial t} &= f_4([E_p], [E-E_p], [N2_a])
\end{aligned}
\tag{6}
$$

with

$$
f_1 = ([E_t]-[E_p]-[E-E_p])(\alpha_1([E_t]-[E_p]-[E-E_p]) + \alpha_2[E_p] + \alpha_3[E-E_p]) -
$$

$$
\gamma_1[RG_a][E_p] - \gamma_2[N2_a][E_p] - k_{on}([EGF_t]-[E-E_p])[E_p]^2 + 1/2 k_{off}[EE_p];
$$

$$
f_2 = k_{on}([EGF_t]-[E-E_p])([E_p]^2 + ([E_t]-[E_p]-[E-E_p])^2) - k_{off}[E-E_p];
$$

$$
f_3 = k_1([RG_t]-[RG_a]) - k_2[RG_a] - \beta_1[RG_a]([E_p]+[E-E_p]);
$$

and

$$
f_4 = \epsilon(k_1([N2_t]-[N2_a]) - k_2[N2_a] + \beta_2([E_p]+[E-E_p])([N2_t]-[N2_a])).
$$

The reaction terms are described in (Stanoev et al., 2018). $[E-E_p]$ is the phosphorylated ligand-bound dimeric EGFR, $[E_p]$ - ligandless phosphorylated EGFR, $[E_t]$ - total amount of EGFR, $[RG_a], [RG_t]$ and $[N2_a], [N2_t]$ - the active and total amount of the membrane localized PTPRG and the ER-bound PTPN2, respectively. Both, the receptor and the deactivating enzymes have active and inactive states, and the model equations describe their state transition rates. Therefore, mass is conserved in the system and the total protein concentrations of the three species ($[E_t]$, $[RG_t]$ and $[N2_t]$) are constant parameters. Autonomous, autocatalytic and ligand-bound-induced activation of ligandless EGFR ensue from bimolecular interactions with distinct rate constants $\alpha_{1-3}$, respectively. Other parameters are as follows: $k_1/k_2$ — activation/inactivation rate constants of the phosphatases, $\beta_1/\beta_2$ - receptor-induced regulation rate constants of $PTPRG/PTPN2$, $\gamma_1/\gamma_2$ - specific reactivity of the enzymes ($PTPRG/PTPN2$) towards the receptor. The EGFR-PTPN2 negative feedback is on a time scale ($\epsilon$) approximately two orders of magnitude slower than the phosphorylation-dephosphorylation reaction, as estimated from the $\sim 4min$ recycling time of $EGFR_p$ (Stanoev et al., 2018). This enables, when necessary, to consider a quasi-steady state approximation for the dynamics of PTPN2 for simplicity:

$$[N2_a]_{qss} = [N2_t] \cdot \frac{(k_1 + \beta_2 \cdot ([E_p] + [E - E_p]))}{k_1 + k_2 + \beta_2 \cdot ([E_p] + [E - E_p])} \tag{7}$$

$[EGF_t]$ denotes the total ligand concentration. Assuming that at low, physiologically relevant EGF doses, the ligand will be depleted from the solution due to binding to EGFR (Lauffenburger and Linderman, 1996), ligand-binding unbinding was explicitly modeled ($k_{on}$, $k_{off}$) in Eqs.6.

The diffusion terms model the lateral diffusion of the EGFR and PTPRG molecules on the plasma membrane, whereas PTPN2 is ER-bound and does not diffuse. Single particle tracking studies have demonstrated that EGFR molecules on the plasma membrane occupy three distinct mobility states, free, confined and immobile, with the occupations of the free and immobile states decreasing and increasing significantly after EGF stimulation ($2min$ after EGF stimulation, corresponding with the time-scale of EGF binding) (Ibach et al., 2015). In the reaction-diffusion (RD) simulations therefore for simplicity, it is assumed that $D_{E-E_p} \approx 0$, whereas diffusion constants of same order are assumed for the ligandless EGFR and PTPRG ($D_{E_p} \sim D_{RG_a}$).

#### 0.3 Analytical consideration for an $SN_{PB}$ existence in the EGFR network

To identify analytically the existence of a $SN_{PB}$ in the EGFR receptor network, we performed a weakly nonlinear analysis as described in the general consideration (Section 0.1). For this, we considered the system Eqs.(6), where the dynamics of PTPN2 is at quasi-steady state (Eq.(7)), $[E - E_p] = 0$, and rest of the dependent and independent variables were scaled to have a dimensionless form. Let $[\tilde{E_p}] = [E_p]/E_0$, $[\tilde{RG_a}] = [RG_a]/RG_0$, $\tilde{x} = x/x_0$, $\tau = t/t_0$, such that $t_0 = 1/(k_1 + k_2)$, $E_0 = k_1/\beta_2$, $RG_0 = (k_1 + k_2)/\gamma_1$ and $t_0/x_0^2 = 1/D_{E_p}$. Substituting these into Eqs.(6) yields the system of dimensionless equations:

$$\begin{aligned} \frac{\partial[\tilde{E_p}]}{\partial\tau} &= q_1 + q_2[\tilde{E_p}] + q_3[\tilde{E_p}]^2 - [\tilde{RG_a}][\tilde{E_p}] - \frac{q_4(1 + [\tilde{E_p}])[\tilde{E_p}]}{(1 + k + [\tilde{E_p}])} + \frac{\partial^2[\tilde{E_p}]}{\partial\tilde{x}^2} \\ \frac{\partial[\tilde{RG_a}]}{\partial\tau} &= r_1 - [\tilde{RG_a}] - r_2[\tilde{RG_a}][\tilde{E_p}] + D\frac{\partial^2[\tilde{RG_a}]}{\partial\tilde{x}^2} \end{aligned} \tag{8}$$

with $q_1 = \frac{a_1 \cdot [E_t]^2 \cdot k_3}{(k_1+k_2)\cdot\beta_2}$, $q_2 = \frac{(a_2 - 2\cdot a_1)\cdot[E_t]}{k_1+k_2}$, $q_3 = \frac{(a_1-a_2)\cdot k_1}{(k_1+k_2)\cdot\beta_2}$, $q_4 = \frac{\gamma_2\cdot[N2_t]}{k_1+k_2}$, $k = k_2/k_1$, $r_1 = \frac{k_1\cdot[RG_t]\cdot\gamma_1}{(k_1+k_2)^2}$, $r_2 = \frac{\beta_1\cdot k_1}{(k_1+k_2)\cdot\beta_2}$ and $D = \frac{D_{RG_a}}{D_{E_p}}$.

We further simplify the system Eqs.(8) by taking the Talyor series expansion of the quasi-steady state approximation of $[N2_a]$ around $E_s$, the steady state of $[\tilde{E_p}]$:

$$\frac{q_4(1 + [\tilde{E_p}])[\tilde{E_p}]}{1 + k + [\tilde{E_p}]} = q_7 + q_8[\tilde{E_p}] + q_9[\tilde{E_p}]^2 + o([\tilde{E_p}]^2) \tag{9}$$

with $q_7 = \frac{E_s q_4}{1+k+E_s} - \frac{E_s q_4(1+k)}{(1+k+E_s)^2}$, $q_8 = \frac{E_s q_4}{1+k+E_s} + \frac{q_4(1+k)}{(1+k+E_s)^2}(1 - E_s)$ and $q_9 = \frac{q_4(1+k)}{(1+k+E_s)^2}$, thus yielding:

$$
\begin{aligned}
\frac{\partial[\tilde{E}_p]}{\partial\tau} &= q_9 + q_{10}[\tilde{E}_p] + q_{11}[\tilde{E}_p]^2 - [\widetilde{RG}_a][\tilde{E}_p] + \frac{\partial^2[\tilde{E}_p]}{\partial\tilde{x}^2} \\
\frac{\partial[\widetilde{RG}_a]}{\partial\tau} &= r_1 - [\widetilde{RG}_a] - r_2[\widetilde{RG}_a][\tilde{E}_p] + D\frac{\partial^2[\widetilde{RG}_a]}{\partial\tilde{x}^2}
\end{aligned}
\tag{10}
$$

with $q_9 = q_1 - q_7$, $q_{10} = q_2 - q_8$ and $q_{11} = q_3 - q_9$.

To avoid long expression in the further analysis, we re-name the dependent variables as $u_1 = [\tilde{E}_p]$ and $u_2 = [\widetilde{RG}_a]$, and the independent variables as $\tilde{x} = x$, $\tau = t$ . The system Eqs.(8) therefore obtains the generic form:

$$
\begin{aligned}
\frac{\partial u_1}{\partial t} &= F_1(u_1, u_2) + \frac{\partial^2 u_1}{\partial x^2} \\
\frac{\partial u_2}{\partial t} &= F_2(u_1, u_2) + D\frac{\partial^2 u_2}{\partial x^2}.
\end{aligned}
\tag{11}
$$

In order to perform linear stability analysis, a one-dimensional projection of Eq.(11) is considered,

$$
\begin{aligned}
\frac{du_{1L}}{dt} &= F_1(u_{1L}, u_{2L}) - (u_{1L} - u_{1R}) = G_1(u_{1L}, u_{2L}, u_{1R}) \\
\frac{du_{2L}}{dt} &= F_2(u_{1L}, u_{2L}) - D(u_{2L} - u_{2R}) = G_2(u_{1L}, u_{2L}, u_{2R}) \\
\frac{du_{1R}}{dt} &= F_1(u_{1R}, u_{2R}) - (u_{1R} - u_{1L}) = G_3(u_{1R}, u_{2R}, u_{1L}) \\
\frac{du_{2R}}{dt} &= F_2(u_{1R}, u_{2R}) - D(u_{2R} - u_{2L}) = G_4(u_{1R}, u_{2R}, u_{2L})
\end{aligned}
\tag{12}
$$

Let, $\mathbf{U_s} = \begin{pmatrix} u_{1Ls} \\ u_{2Ls} \\ u_{1Rs} \\ u_{2Rs} \end{pmatrix}$ be the stable symmetric steady state of the system ($u_{1Ls} = u_{1Rs}$, $u_{2Ls} = u_{2Rs}$). A small amplitude perturbation on this symmetric steady state of the form,

$$\begin{pmatrix} u_{1L}(t) \\ u_{2L}(t) \\ u_{1R}(t) \\ u_{2R}(t) \end{pmatrix} = \begin{pmatrix} u_{1Ls} \\ u_{2Ls} \\ u_{1Rs} \\ u_{2Rs} \end{pmatrix} + \begin{pmatrix} \delta u_{1L} \\ \delta u_{2L} \\ \delta u_{1R} \\ \delta u_{2R} \end{pmatrix} \cdot e^{\lambda t} \tag{13}$$

yields a linearized equation,

$$\lambda \begin{pmatrix} \dfrac{d\delta u_{1L}}{dt} \\ \dfrac{d\delta u_{2L}}{dt} \\ \dfrac{d\delta u_{1R}}{dt} \\ \dfrac{d\delta u_{2R}}{dt} \end{pmatrix} = \mathbf{J} \begin{pmatrix} \delta u_{1L} \\ \delta u_{2L} \\ \delta u_{1R} \\ \delta u_{2R} \end{pmatrix} \tag{14}$$

where $\mathbf{J} = \begin{pmatrix} \frac{\partial G_1}{\partial u_{1L}} & \frac{\partial G_1}{\partial u_{2L}} & \frac{\partial G_1}{\partial u_{1R}} & 0 \\ \frac{\partial G_2}{\partial u_{1L}} & \frac{\partial G_2}{\partial u_{2L}} & 0 & \frac{\partial G_2}{\partial u_{2R}} \\ \frac{\partial G_3}{\partial u_{1L}} & 0 & \frac{\partial G_3}{\partial u_{1R}} & \frac{\partial G_3}{\partial u_{2R}} \\ 0 & \frac{\partial G_4}{\partial u_{2L}} & \frac{\partial G_4}{\partial u_{1R}} & \frac{\partial G_4}{\partial u_{2R}} \end{pmatrix}$

is the Jacobian of the system evaluated at the symmetric steady state. In order to identify existence of PB in the system, the condition given in Eq.(3) should be satisfied for an odd mode of the perturbation. For the one-dimensional projection (Eqs.(12)), the odd mode of the perturbation $(\delta\mathbf{U}(-\mathbf{x})) = -\delta\mathbf{U}(\mathbf{x}))$ must yield: $\delta u_{1L} = -\delta u_{1R}$ and $\delta u_{2L} = -\delta u_{2R}$. Substituting this into Eq.(14) to obtain $F_{-}(\lambda)$, in the limit $\lambda \to 0$ renders:

$$\lim_{\lambda\to 0} F_{-}(\lambda) = det \begin{pmatrix} (\frac{\partial G_1}{\partial u_{1L}} + \frac{\partial G_3}{\partial u_{1R}}) - (\frac{\partial G_1}{\partial u_{1R}} + \frac{\partial G_3}{\partial u_{1L}}) & (\frac{\partial G_1}{\partial u_{2L}} + \frac{\partial G_2}{\partial u_{2R}}) \\ (\frac{\partial G_2}{\partial u_{1L}} + \frac{\partial G_4}{\partial u_{1R}}) & (\frac{\partial G_2}{\partial u_{2L}} + \frac{\partial G_4}{\partial u_{2R}}) - (\frac{\partial G_2}{\partial u_{2R}} + \frac{\partial G_4}{\partial u_{2L}}) \end{pmatrix} = 0 \tag{15}$$

Thus, there exists parameter set for which existence of PB in the system Eq.(12) is guaranteed.

To identify whether the PB is sub-critical and thereby identify existence of a $SN_{PB}$, the solution of the system Eqs.(11) is approximated as in Eq.(4):

$$
\begin{aligned}
u(x,t) &= \phi(t)e^{ik_m x} + \phi^*(t)e^{-ik_m x} + u_0(t) + \sum_{n=2}^{3}(u_n(t)e^{nik_m x} + u_n^*(t)e^{-nik_m x}) \\
v(x,t) &= \phi(t)e^{ik_m x} + \phi^*(t)e^{-ik_m x} + v_0(t) + \sum_{n=2}^{3}(v_n(t)e^{nik_m x} + v_n^*(t)e^{-nik_m x})
\end{aligned}
\tag{16}
$$

The expansion is taken to $n = 3^{rd}$ order, rendering an amplitude equation of $5^{th}$ order. As described in (Becherer et al., 2009), the complex coefficients of the $n = 0^{th}, n = 2^{nd}$ and $n = 3^{rd}$ harmonics can be approximated as power series of $\phi(t)$. Substituting into Eqs.(11) allows to derive these coefficients. This yields a system of coupled ODEs representing the time evolution of the complex amplitudes, in this case, for $\phi(t)$, $u_0(t)$, $v_0(t)$, $u_1(t)$, $v_1(t)$, $u_2(t)$, $v_2(t)$, $u_3(t)$ and $v_3(t)$. Assuming that the dynamics of the higher order harmonics reaches their steady state much faster than the leading perturbation does, the derivatives of their amplitudes can be set to zero. This allows to obtain expressions of the amplitudes purely as functions $\phi$ and the parameters of the system as:

$$
\begin{aligned}
u_0(\phi) &= (\frac{1}{q_{10}}(2(1-q_{11}) - \frac{q_9}{|\phi|^2}))|\phi|^2 \\
v_0(\phi) &= (\frac{r_1}{|\phi|^2} - 2r_2)|\phi|^2 \\
u_2(\phi) &= u_2^{(2)}\phi^2 \\
v_2(\phi) &= v_2^{(2)}\phi^2 \\
u_3(\phi) &= u_3^{(3)}\phi^3 \\
v_3(\phi) &= v_3^{(3)}\phi^3
\end{aligned}
\tag{17}
$$

where $u_2^{(2)} = \frac{1-q_{11}}{q_{10}-4k_m^2}$, $v_2^{(2)} = \frac{-r_2}{1+4Dk_m^2}$, $u_3^{(3)} = \frac{u_2^{(2)}+v_2^{(2)}-2q_{11}u_2^{(2)}}{q_{10}-9k_m^2}$ and $v_3^{(3)} = \frac{-r_2(u_2^{(2)}+v_2^{(2)})}{1+9Dk_m^2}$. The dynamics of the leading harmonics ($n = 1$) can be written as:

$$\frac{d\phi}{dt} = c_1\phi + c_2\phi^3 - c_3\phi^5 \qquad (18)$$

where $c_1 = q_{10} - k_m^2 - r_1 + \frac{q_9(1-2q_{11})}{q_{10}}$, $c_2 = (1-q_{11})(2q_{11}-1)(\frac{2}{q_{10}} - \frac{1}{q_{10}-4k_m^2}) + r_2(2 + \frac{1}{1+4Dk_m^2})$ and $c_3 = 2q_{11}u_2^{(2)}u_3^{(3)} - u_2^{(2)}v_3^{(3)}$. Eq.(18) is of Stuart-Landau type and represents a normal form of a sub-critical pitchfork bifurcation. This shows the existence of $SN_{PB}$ in the EGFR network.

To corroborate this, we also performed numerical bifurcation analysis on one-dimensional projection (Eqs.(12)) where the reaction terms have the form as defined in Eqs.(6), including the full form for $[N2_a]$, when $[E - E_p] = 0$. The bifurcation analysis (Figure S1B) was obtained using the Xppaut software package (Ermentrout, 2016). The parameters in the model Eqs.(6) have been described in (Stanoev et al., 2018), where they were calibrated with experimental data: $\alpha_1 = 0.001$, $\alpha_2 = 0.3$, $\alpha_3 = 0.7$, $\beta_1 = 11$, $\beta_2 = 1.1$, $k_1 = 0.5$, $k_2 = 0.5$, $g_1 = 1.9$, $g_2 = 0.1$, $k_{on} = 0.05$, $k_{off} = 0.28$, $\epsilon = 0.01$, $RG_t = 1$, $N2_t = 1$; and the diffusion-like terms have been scaled from the values derived in (Orr et al., 2005): $D_{RG_a} = 0.02$, $D_{E_p} = 0.02$.

The bifurcation analysis demonstrates that for the spatially-distributed EGFR network, the homogeneous steady state (HSS, grey solid line, Figure S1B) losses stability via a symmetry-breaking pitchfork bifurcation (PB), giving rise to inhomogeneous steady states (IHSS), stabilised via saddle-node bifurcations ($SN_{PB}$) (Figure S1B, magenta branched lines). There is a coexistence between the HSS and the IHSS before the PB, rendering it sub-critical. The IHSS (Koseska et al., 2013) is a single attractor that describes a heterogeneous state with two branches corresponding to orientation of the front-back-polarized state. The described IHSS solution is therefore fundamentally distinct from a bistable system where the high and the low phosphorylation states correspond to two different homogeneous steady states. As the IHSS is a single attractor, the high and low phopshorylation state are interdependent, rendering the PB a unique mechanism for generating robust front-back polarization.

The reaction diffusion simulations were performed by assuming PTPN2 at quasi-steady state. The cell boundary was represented with a 1D circular domain of length $L = 2\pi R$ (where $R = 2\mu m$ ) which was then divided into 20 equal bins. The diffusion terms were approximated by central difference method, enabling for conversion of the PDE system to a system of ordinary differential equations (ODEs). Stochastic simulations with additive white noise were implemented by adding $\sigma \cdot dW_t$ ( $\sigma = 0.02$, $dW_t$ is sampled from a normal distribution with mean 0 and variance 0.01) in the equation for $[E_p]$. The stochastic *sdeint* Python package was used. Parameters: $D_{E_p} = D_{RG_a} = 0.008\ \mu m^2/min$. $D_{E_p}$ was taken from (Orr et al., 2005) and scaled to correspond to a cell with perimeter $L$ in the simulations. For organization in the homogenous symmetric steady states (the basal and pre-activated states), organization at criticality or in the asymmetric IHSS, $E_t \in \{0.9, 1.8, 1.255, 1.32\}$ respectively, time step was set to $0.01min$, other parameters as above. Periodic boundary conditions were used. To mimic the dynamic nature of $EGF^{647}$ gradient, a Gaussian function on a periodic window with varying amplitude and standard deviation was used (shape shown in Figure 1D, top). To represent the state-space trajectory (Figure 1F, bottom), stochastic realization of the one-dimensional projection of the full system (as for the bifurcation analysis) was used.

#### 0.4 Physical model of single-cell chemotaxis

To describe signal-induced cell shape changes and subsequent cell migration, we combined the dynamical description of the gradient sensing capability of the EGFR network (Eqs.6, Figure 1B) together with a physical model for cellular migration, thereby implicitly modeling the signal-induced cell shape changes (Figure 1C). In order to couple a mechanical model of the cell with the biochemical EGFR signaling model as a means to simulate large cellular deformations, we utilized the Level Set Method (LSM) (Osher and Sethian, 1988) as described in (Yang et al., 2008). Briefly, the cell boundary at time $t$ is described on a two-dimensional Cartesian grid by the closed-contour $\Gamma(t) = \{\mathbf{x}|\Psi(\mathbf{x},t) = 0\}$, that represent the zero-level set of the potential function $\Psi(\mathbf{x},t)$, taken to have an initial form:

$$\Psi(\mathbf{x},0) = \begin{cases} -d(\mathbf{x},\Gamma), & \text{if } \mathbf{x} \in S \\ d(\mathbf{x},\Gamma), & \text{if } \mathbf{x} \notin S \\ 0, & \text{if } \mathbf{x} \in \Gamma \end{cases} \tag{19}$$

where $S$ identifies the area occupied by the cell and $d(\mathbf{x},\mathbf{\Gamma})$ is the distance of position $x$ to the curve $\Gamma$. Thus, the cell membrane is represented implicitly through the potential function which is defined on the fixed Cartesian grid, eliminating the need to parameterize the boundary, and thereby enabling to handle complex cell boundary geometries.

The shape of the cell ($\Gamma(\mathbf{x},t)$) evolves according to the Hamilton-Jacobi equation:

$$\frac{\partial\Psi(\mathbf{x},t)}{\partial t} + \mathbf{v}(\mathbf{x},t)\cdot\nabla\Psi(\mathbf{x},t) = 0 \tag{20}$$

The vector $\mathbf{v}(\mathbf{x},t)$ is the velocity of the level set moving in the outward direction, thereby intrinsically describing the cell's membrane protrusion and retraction velocities that are driven by internally generated mechanical forces (e.g. actin polymerization or myosin-II retraction (Bray, 2000)). To determine how these forces translate to membrane velocity, a mechanical model that describes the viscoelastic behavior of the cell represented as a viscoelastic cortex surrounding a viscous core, is implemented. Following (Yang et al., 2008), the cortex connecting the cell membrane and the cytoplasm is represented by a Voigt model (parallel connection of an elastic element $k_c$ and a viscous element $\tau_c$, whereas the cytoplasm is modeled as a purely viscous element, $\tau_a$, which is placed in series with the Voigt model.

Let $\mathbf{l}(\mathbf{x},t)$, $\mathbf{x} \in \Gamma(t)$ be the viscoelastic state of the cell at time $t$ and at a position $\mathbf{x}$ on the membrane, such that $|\mathbf{l}|$ represents the length of the numerous parallel unconnected spring-damper systems. The viscoelastic state of the cell then evolves according to:

$$\frac{-k_c}{\tau_c}\mathbf{l}(t) + \frac{1}{\tau_c}\mathbf{P_{total}}(t) = \nabla\mathbf{l} \cdot \mathbf{v}(\mathbf{t}) + \frac{\partial \mathbf{l}(t)}{\partial t} \tag{21}$$

where $\nabla$ is the gradient operator, the pressure $\mathbf{P_{total}}(t) = \mathbf{P_{prot}}(t) + \mathbf{P_{retr}}(t) + \mathbf{P_{area}}(t) - \mathbf{P_{ten}}(t)$ is sum of the protrusion, retraction, area conservation, and cortical tension pressures, respectively. The EGFR signaling state ($[E_p]$) directly determines the protrusion/retraction pressure, since high/low signaling activity triggers actin polymerization / myosin-II retraction following:

$\mathbf{P_{prot}}(t) = K_{prot}(([E_p](t) - < [E_p](t) >)/([E_p]_{max}(t) - < [E_P](t) >))\mathbf{n}$ and

$\mathbf{P_{ret}}(t) = -K_{retr}((< [E_p] > -[E_p])/(< [E_P] > -[E_p]_{max}))\mathbf{n}$, where $< . >$ denotes mean at the membrane, $K_{prot}$, $K_{retr}$ - proportionality constants. The cell is assumed to be flat with uniform thickness, such that the 2D area ($A(t)$) of the cell is conserved ($\mathbf{P_{area}}(\mathbf{t}) = K_{area}(A(0) - A(t))\mathbf{n}$), $K_{area}$ - proportionality constant. The pressure generated by the cortical tension therefore depends only on the 2D local surface curvature and the 2D equilibrium pressure, rendering the rounding pressure due to cortical tension to be $\mathbf{P_{ten}}(t) = K_{ten}(\kappa(\Gamma) - 1/R)\mathbf{n}$, with $\kappa(x)$ being the local membrane curvature, R - initial cell radius, was set to 2 $\mu m$, and $K_{ten}$ - proportionality constant. The local membrane velocity $\mathbf{v}(\mathbf{x})$, $\mathbf{x} \in \Gamma(t)$ depends both on the viscoelastic nature of the cell and on the effective pressure profile ($\mathbf{P_{total}}(t)$) and is given by,

$$\mathbf{v} = \frac{-k_c}{\tau_c}\mathbf{l} + (\frac{1}{\tau_c} + \frac{1}{\tau_a})\mathbf{P_{total}} \tag{22}$$

For the simulations in Figures 1, 4 and Figure S5, first the stochastic PDEs (Eqs.(6)) are solved and the kymographs of the signalling ($[E_p]$) activity are generated. The viscoelastic state is initialized with zero value on the membrane, $\mathbf{l}(\mathbf{x}, 0) = 0$. At each time point, $\mathbf{P_{total}}$ is estimated, as well as the local membrane velocity using Eq. (22). This velocity is then used to evolve both the viscoelastic state (Eq. (21)) and the potential function (Eq.(19)).

The spatial discretization of these advection equations (Eqs.(21),(22)) was performed using the *upwindENO2* scheme, as described in the Level Set Toolbox (Mitchell, 2007) and was integrated with first order forward Euler method. The time step was set to $0.01min$ and the potential function was solved on a 2D Cartesian grid with spatial discretization of 5 points per $\mu m$. All the codes were custom implemented in Python. Parameters: $k_c = 0.1 \quad nN/\mu m^3$, $\tau_c = 0.08 \quad nNmin/\mu m^3$, $\tau_a = 0.1 \quad nNmin/\mu m^3$, $K_{prot} = 0.08 \quad nN/\mu m^2$, $K_{retr} = 0.05 \quad nN/\mu m^2$, $K_{area} = 0.02 \quad nN/\mu m^4$, $K_{ten} = 0.1 \quad nN/\mu m$. $K_{ten}$ was taken from the literature, corresponding to an experimentally measured range of cell cortical tension values (Cartagena-Rivera et al., 2016). The rest of the parameters were selected to match the cell migration speed during gradient and memory phase, estimated from the experiments (Figure 3A, $v = 0.49 \pm 0.173 \mu m/min$).

#### Supplementary videos

**Movie S1.** Corresponding to Figure 1F. *In silico* temporal evolution of the state-space trajectory of the EGFR sensing system in $E_p$ - $P_{RG}$ - $P_{N2}$ space.

**Movie S2:** Corresponding to Figure 2F. State-space trajectory reconstructed from experimentally obtained temporal EGFR$^{mCitrine}$ phosphorylation profile (1h during and 3h after EGF$^{647}$ gradient duration) of a representative MCF7-EGFR$^{mCitrine}$ cell. $160min$ from the reconstructed state-space trajectory are shown.

**Movie S3:** Corresponding to Figure 3A. Migration trajectory of a representative MCF10A cell subjected for 5h to dynamic EGF$^{647}$ gradient (green) and 9h after gradient wash-out (red).

**Movie S4:** Corresponding to Figure 3G. State-space trajectory reconstructed from experimentally obtained temporal EGFR$^{mCitrine}$ phosphorylation profile of a representative MCF7-EGFR$^{mCitrine}$ cell under Lapatinib treatment. EGFR phosphorylation was quantified 1h during EGF$^{647}$ gradient and 3h after wash-out with 1$\mu$M Lapatinib. $140min$ from the reconstructed state-space trajectory are shown.

**Movie S5:** Corresponding to Figure 3I. Migration trajectory of a representative MCF10A cell subjected for 5h to dynamic EGF$^{647}$ gradient (green) and 9h after gradient wash-out with 3$\mu$M Lapatinib (red).

**Movie S6.** Corresponding to Figure 4B. *In silico* evolution of a cellular response to a dynamic chemical field for organization at criticality. EGFR phosphorylation (blue-to-yellow/low-to-high), cell shape and migration trajectory are shown during (green/orange) and after (red) EGF gradient presence, as obtained from a physical model of single-cell chemotaxis (Supplementary information).

**Movie S7.** Corresponding to Figure S5B, C. *In silico* evolution of a cellular response to a dynamic chemical field for organization in the stable inhomogenous steady state (cell polarization regime). Notations as in Movie S6.

**Movie S8:** Corresponding to Figure 4C. Cellular navigation in a changing gradient field. Migration trajectory of a representative MCF10A cell subjected to a spatial-temporal $EGF^{647}$ gradient field described in Figure 4A.

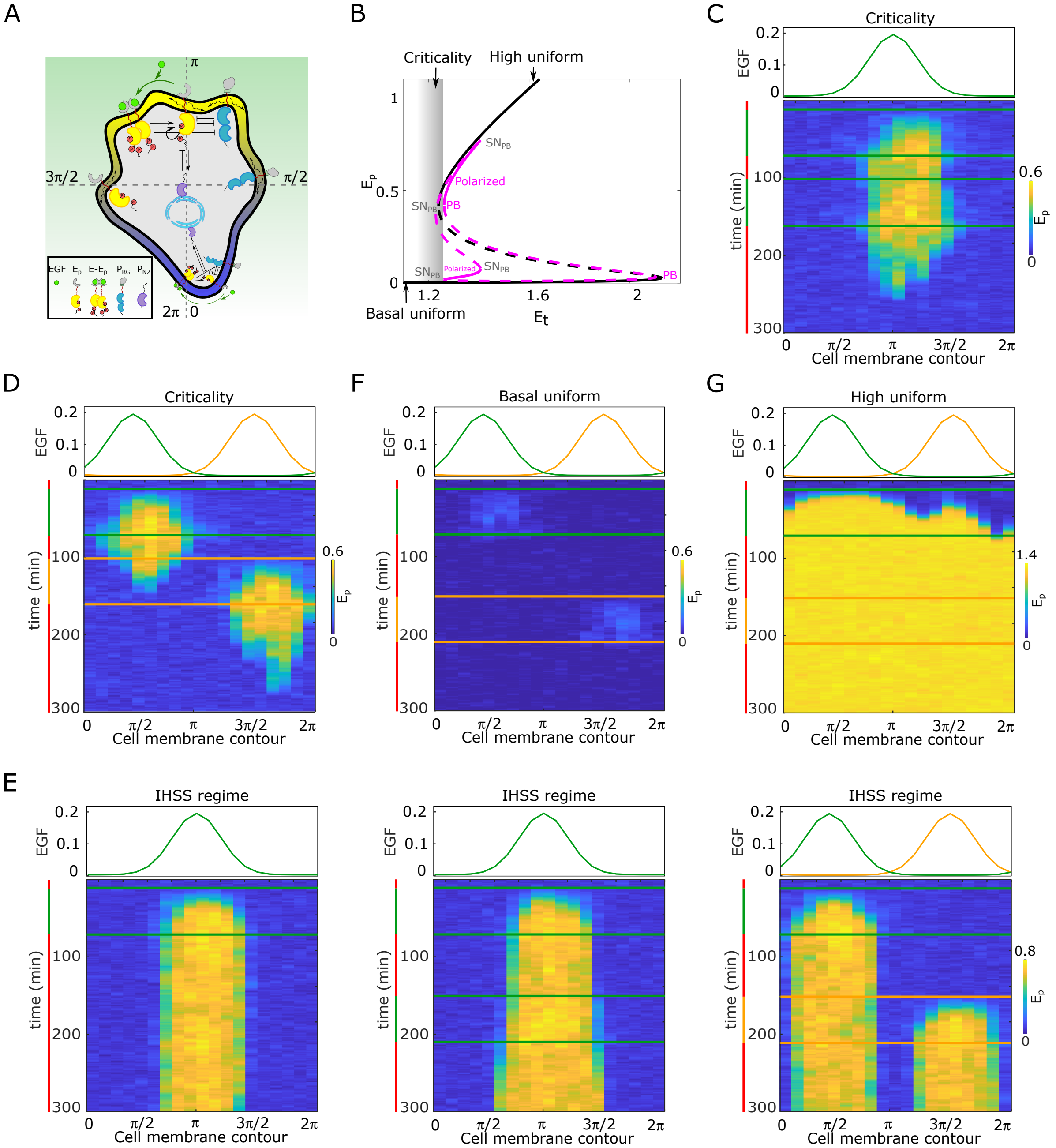
A
B
C
D
F
G
E
Criticality
High uniform
Polarized
PB
Basal uniform
Et
Ep
Cell membrane contour
time (min)
EGF
Basal uniform
High uniform
IHSS regime

**Figure S1, related to Figure 1. Features of receptor activity for different organization in parameter space.** **A**, Spatial representation of the EGFR sensing network shown in Figure 1B, with equivalent notations and line descriptions. **B**, Bifurcation diagram of the EGFR sensing network. Notations and line description as in Figure 1A. $E_t$: total EGFR on the plasma membrane. Parameters in Supplementary information. **C**, Top: Position of two subsequent dynamic EGF gradients in the numerical simulation (profiles as in Figure 1D, top). Bottom: Representative *in silico* kymograph of EGFR phosphorylation ($E_p$) for organization of the system at criticality. Shape changes depicted in Figure 1H, left. **D**, Same as in (**C**), only when the second gradient (yellow) is from the opposite direction. Corresponding shape changes depicted in Figure 1H, right. **E**, Position of dynamic EGF signals(s) in the numerical simulation (top) and respective kymographs of EGFR activity changes (bottom) for organization of the system in the stable inhomogeneous state (magenta attractor in (**B**)). Left: Single dynamic gradient; Middle: a temporally disrupted gradient represented by two subsequent dynamic gradients from the same direction; Right: Second gradient (orange) from the opposite direction. **F**, Same as in (**D**), only for organization in the homogeneous steady state representing symmetric basal EGFR phosphorylation (lower solid black line in **B**). **G**, Same as in **D**, only for organization in the homogeneous steady state representing uniform high EGFR phosphorylation (upper solid black line in **A**). For **B-G**, parameters in Supplementary information. Vertical green(orange)/red lines: stimulus presence/absence.

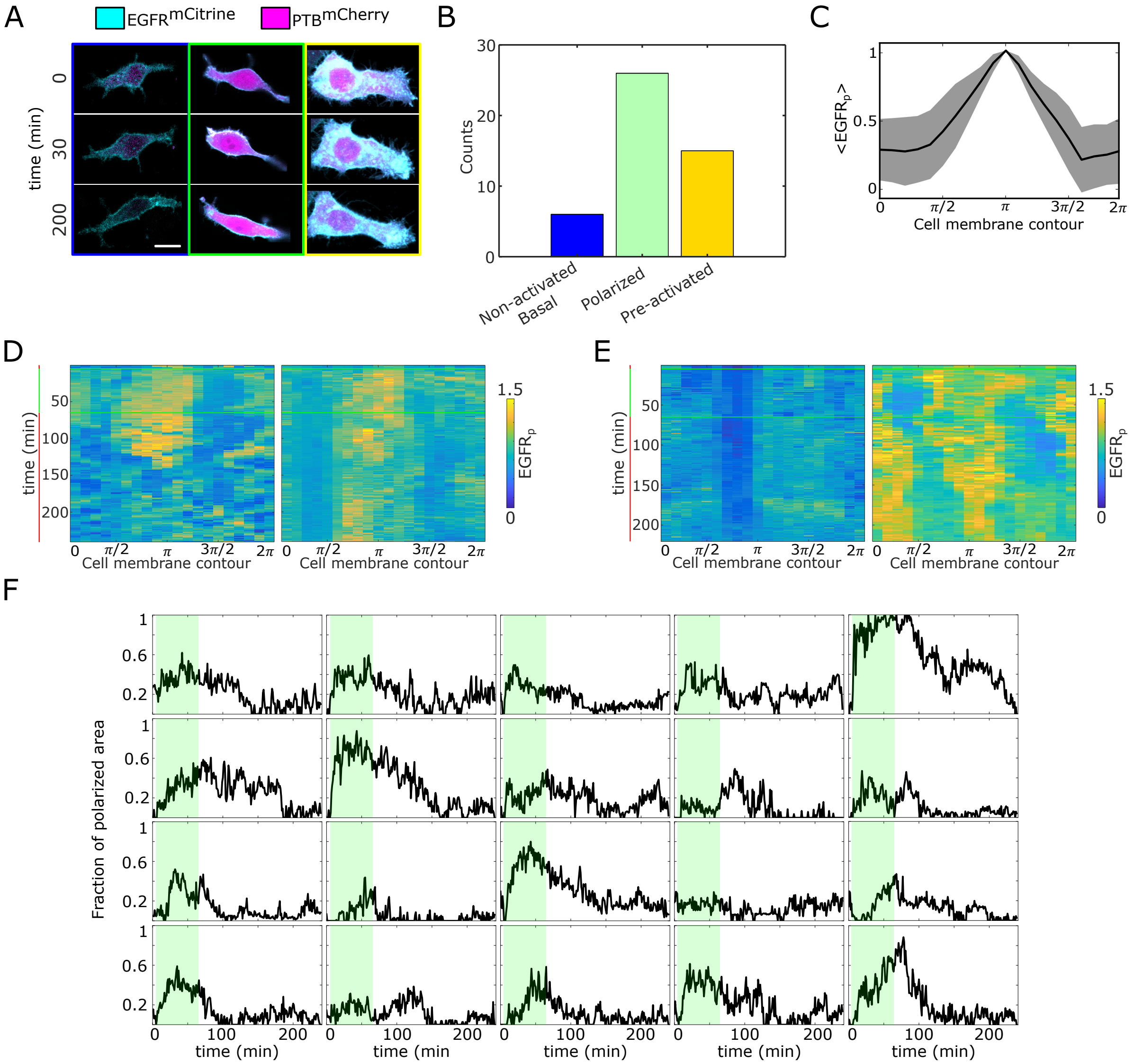
A
EGFR^mCitrine
PTB^mCherry
time (min)
0
30
200
B
Counts
Non-activated Basal
Polarized
Pre-activated
C
<EGFR_p>
Cell membrane contour
D
time (min)
Cell membrane contour
EGFR_p
E
time (min)
Cell membrane contour
EGFR_p
F
Fraction of polarized area
time (min)
time (min

**Figure S2, related to Figure 2. Quantification of EGFR$^{mCitrine}$ phosphorylation polarization.** **A** Representative images / overlay of EGFR$^{mCitrine}$ (cyan) and PTB$^{mCherry}$ (magenta) prior to ($0min$), during ($30min$) and after ($200min$) MCF7-EGFR$^{mCitrine}$ cells were subjected to $60min$ EGF$^{647}$ gradient. Columns: non-activated (blue), transiently polarized (green) and uniformly pre-activated (yellow). Scale bar: $15\mu m$. **B**, Distribution of single-cell responses corresponding to **A**, $N = 7$. **C**, Average profile of the spatial projection of the fraction of phosphorylated EGFR$^{mCitrine}$ from single-cell kymographs. For each cell, temporal average per spatial bin is calculated, and the final spatial profile was estimated as an average of a moving window of 7 points. Peaks of the single-cell distributions were shifted to $\pi$ before averaging. Mean±s.d. from n=20, N=7 is shown. **D**, Additional exemplary single-cell kymographs depicting polarized EGFR$^{mCitrine}$ phosphorylation. Data acquisition and quantification as in Figure 2C. **E**, Same as in **D**, only for non-activated (basal, left) and uniformly pre-acivated (right) EGFR$^{mCitrine}$ phosphorylation. **F**, Temporal profiles of the estimated fraction of polarized area for single cells. Green shaded area: EGF$^{647}$ gradient duration. The mean±s.d. shown in Figure 2D.

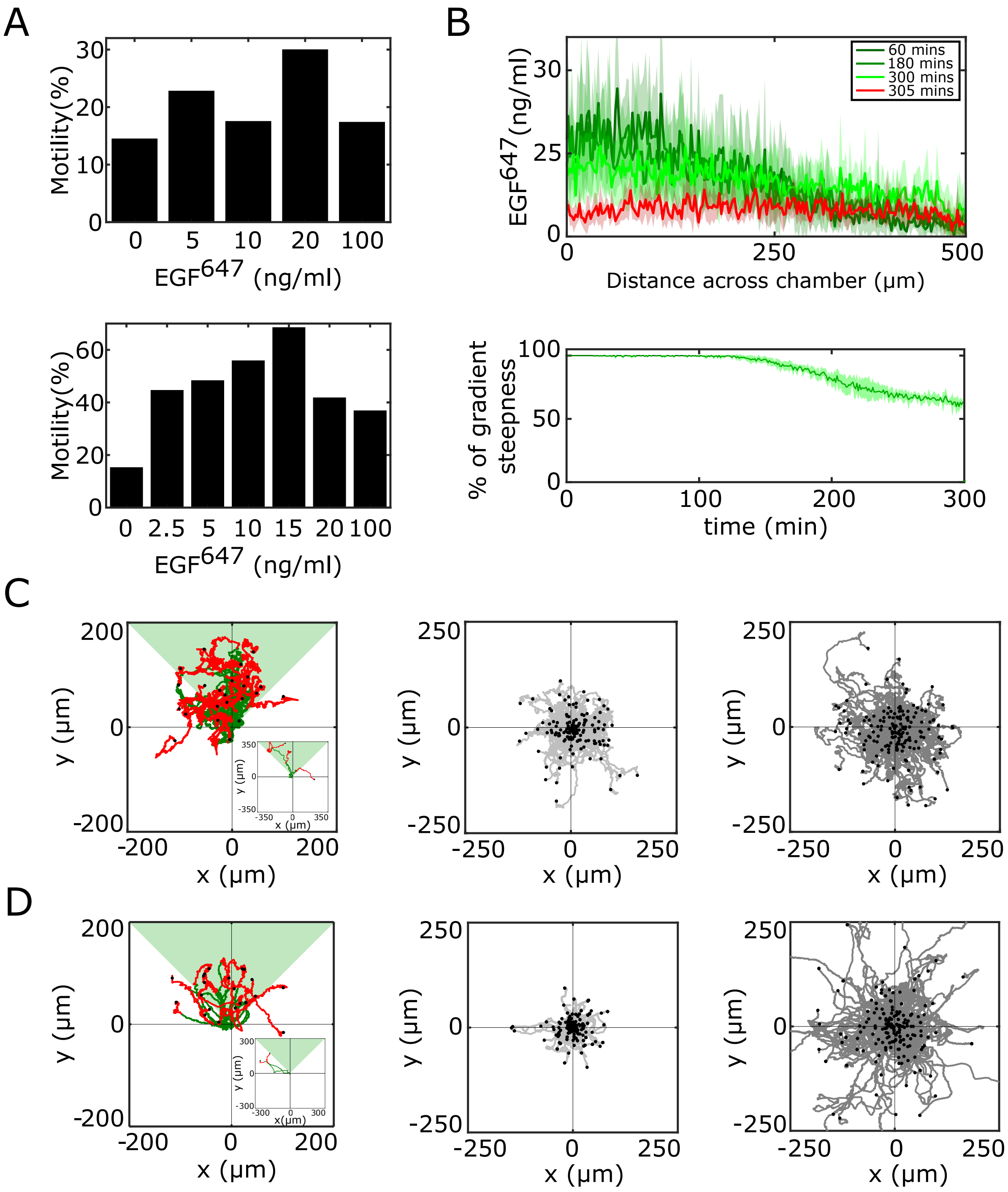
A
Motility(%)
EGF647 (ng/ml)
0 5 10 20 100
0 2.5 5 10 15 20 100
B
EGF647(ng/ml)
60 mins
180 mins
300 mins
305 mins
Distance across chamber (μm)
% of gradient steepness
time (min)
C
D
y (μm)
x (μm)

**Figure S3, related to Figure 3. Characterization of MCF7-EGFR$^{mCitrine}$ and MCF10A single-cell migration.** **A**, Identification of optimal EGF$^{647}$ dose range for single-cell gradient migration assay for MCF7-EGFR$^{mCitrine}$ (top) and MCF10A (bottom). Percentage of cell having motility greater than a displacement threshold ((Number of cell tracks with track length greater than threshold/Total number of cells)*100) is shown. **B**, Top: Quantification of 5h dynamic EGF$^{647}$ gradient at distinct time-points. Bottom: Corresponding quantification of the temporal evolution of the gradient slope. Percentage of gradient steepness: $((EGF^{647}_{(0)} - EGF^{647}_{(L)})/EGF^{647}_{(0)}) * 100$ where $L$ is the length across the chamber. Mean±s.d. from N=4 is shown. **C**, Divergence plots depicting MCF7-EGFR$^{mCitrine}$ single-cell trajectories quantified, left: 5h during (green) and for 9h after (red) dynamic EGF$^{647}$ gradient duration (n=44, N=7); middle: 14h of 0ng/ml EGF$^{647}$ (n=207, N=2); and right: 14h of uniform 20ng/ml EGF$^{647}$ stimulation (n=200, N=2). **D**, Same as in **C**, only for MCF10A cells. Left: n=23, N=5; middle: n=249, N=3; right: n=299, N=3. Related to Figures 3A-C. Black dots: end of tracks.

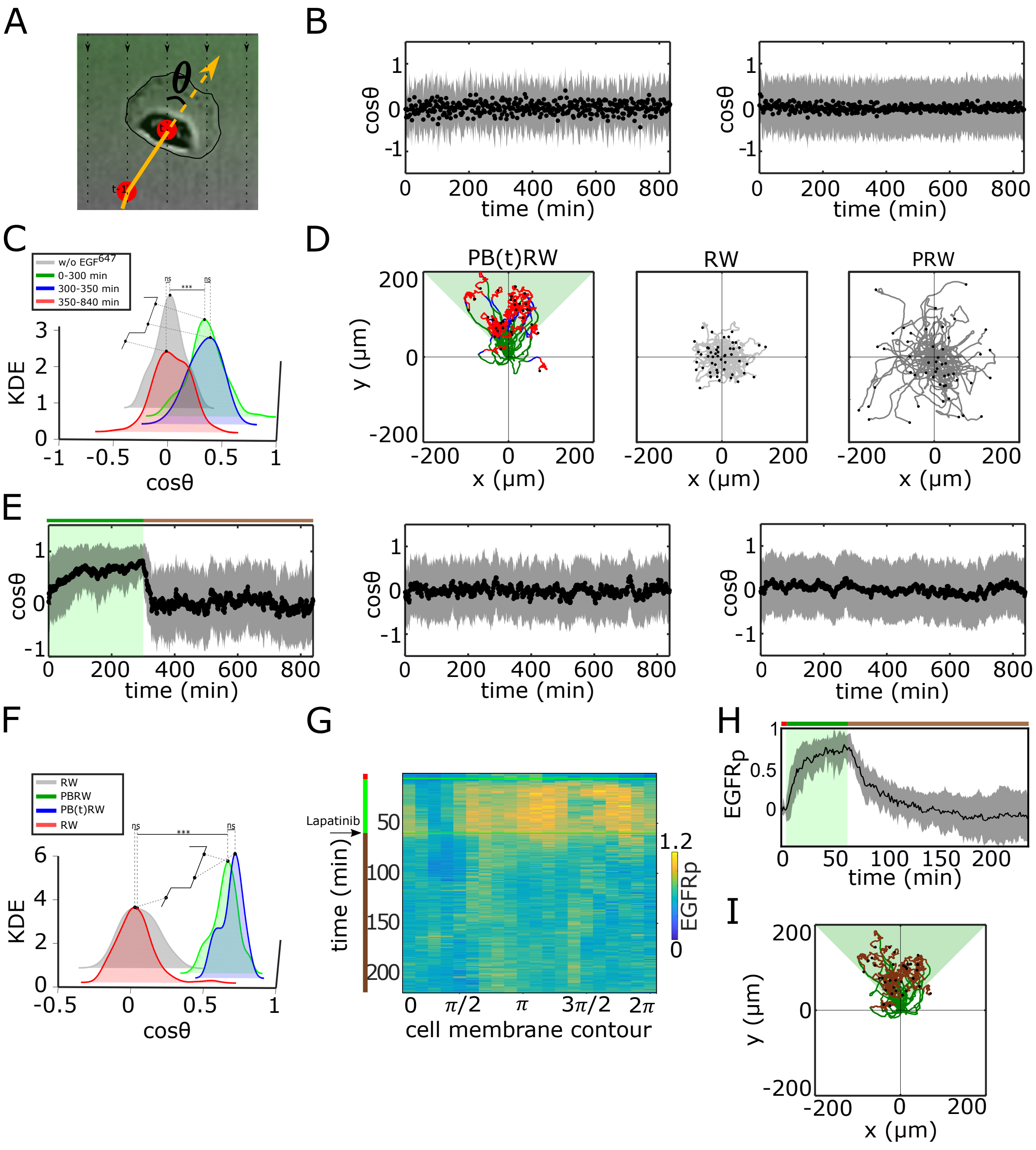
A
θ
B
cosθ
time (min)
C
w/o EGF647
0-300 min
300-350 min
350-840 min
KDE
cosθ
D
PB(t)RW
RW
PRW
y (μm)
x (μm)
E
cosθ
time (min)
F
RW
PBRW
PB(t)RW
RW
KDE
cosθ
G
Lapatinib
time (min)
cell membrane contour
EGFRp
H
EGFRp
time (min)
I
y (μm)
x (μm)

**Figure S4, related to Figure 3. Characterization of single cell migration patterns. A**, Scheme of single-cell turning angle estimation ($\cos\theta$). **B**, Average $\cos\theta$ from single MCF10A cell trajectories (mean±sd), estimated over a $2min$ interval upon, left: 0ng/ml EGF$^{647}$ (n=249, N=3); right: 20ng/ml uniform EGF$^{647}$ stimulation (n=299, N=3). Related to Figure 3A-C. **C**, Kernel density estimates (KDE) of the distributions in (**B**) and Figure 3C top, in continuous EGF$^{647}$ absence (grey), during 5h dynamic EGF$^{647}$ gradient (green), after gradient wash-out: $t \in [300min, 350min]$ (blue) and $t \in [350min, 840min]$ (red). p-values: $***$, p≤0.001, ns: not significant, KS-test. **D**, Synthetic single-cell trajectories (Eq. (6), Methods). Left: Persistent biased random walk PB(t)RW; middle: random walk (RW); right: Persistent random walk (PRW). Parameters: for PB(t)RW, $\tau = 38.143$, $b(t) = 0.134$, $D = 2.207$ for $t \in [0min, 350min]$ (green,blue), $\tau = 11.105$, $b(t) = 0$, $D = 0.425$ for $t \in [350min, 840min]$ (red); for RW, $\tau = 11.105$, $b(t) = 0$, $D = 0.425$; for PRW, $\tau = 38.143$ and $D = 2.207$. **E**, Same as in **B.**, only from the synthetic trajectories. Left: PB(t)RW with $\tau = 38.143$, $D = 2.207$, $b(t) = 0.134$ for $t \in [0min, 300min]$ (green shading), $\tau = 11.105$, $D = 0.425$, $b(t) = 0$ for $t \in [300min, 840min]$, middle: RW; right: PRW. **F**, Same as in **C**, only from the synthetic trajectories. **G**, Exemplary single-cell kymograph depicting phosphorylated EGFR$^{mCitrine}$ for data acquired at $1min$ intervals in live MCF7-EGFR$^{mCitrine}$ cell subjected to $60min$ EGF$^{647}$ gradient, and 3h after gradient wash-out with 1 µM Lapatinib. **H**, Average temporal profiles of plasma membrane EGFR$^{mCitrine}$ phosphorylation of live MCF7-EGFR$^{mCitrine}$ cells subjected to $60min$ EGF$^{647}$ gradient, and 3h after gradient was-out with 1 µM Lapatinib. Related to Figure 3G. Mean±s.d. from n=9, N=2 is shown. Green shaded area: EGF$^{647}$ gradient. **I**, Synthetic single cell trajectories generated when PBRW is considered only in the time frame during gradient duration mimic the experimental data in Figure 3I. Parameters as in (**E**, left).

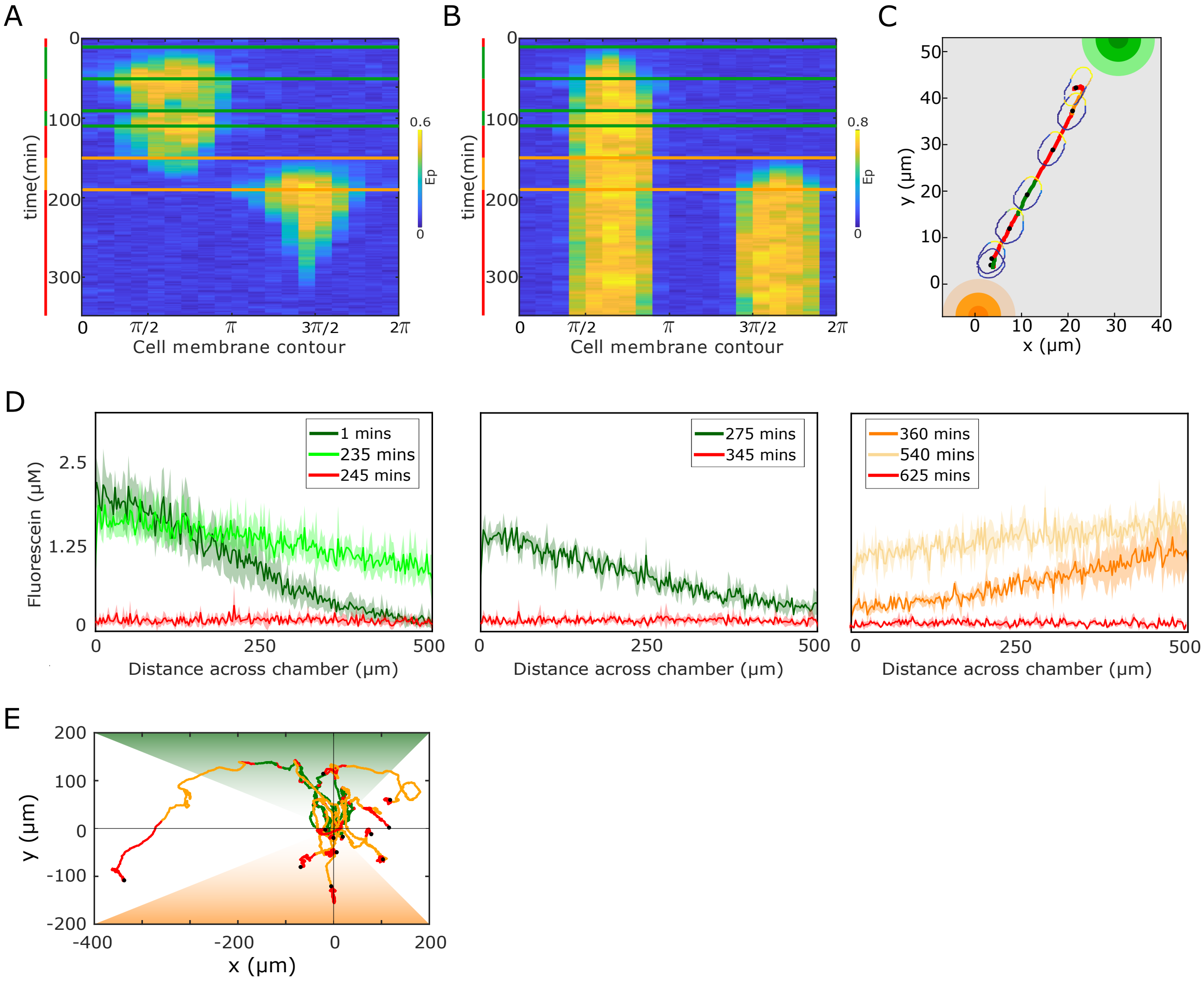


**Figure S5, related to Figure 4. Single-cell navigation in changing growth factor fields. A**, *In silico* obtained $E_p$ kymograph corresponding to Figure 4B. **B**, $E_p$ kymograph obtained for organization in the stable inhomogeneous steady state, when a cell is subjected to the gradient filed in Figure 4A. **C**, Corresponding changes in cellular morphology and respective motility trajectory over time. Trajectory and $E_p$ color coding as in Figure 4A. Cell size is magnified for better visibility. See also movie S7. **D**, Quantification of a 15h dynamic $EGF^{647}$ gradient field at distinct time-points. Mean±s.d. from N=2 is shown. **C**, Divergence plots depicting MCF10A single-cell trajectories quantified during migration in dynamic $EGF^{647}$ gradient filed shown in (**D**). n=12, N=5.
